# Control of neuronal excitation-inhibition balance by BMP-SMAD1 signaling

**DOI:** 10.1101/2023.03.11.532164

**Authors:** Zeynep Okur, Nadia Schlauri, Vassilis Bitsikas, Myrto Panopoulou, Raul Ortiz, Michaela Schwaiger, Kajari Karmakar, Dietmar Schreiner, Peter Scheiffele

## Abstract

Throughout life, neuronal networks in the mammalian neocortex maintain a balance of excitation and inhibition which is essential for neuronal computation. Deviations from a balanced state have been linked to neurodevelopmental disorders and severe disruptions result in epilepsy. To maintain balance, neuronal microcircuits composed of excitatory and inhibitory neurons sense alterations in neural activity and adjust neuronal connectivity and function. Here, we identified a signaling pathway in the adult mouse neocortex that is activated in response to elevated neuronal network activity. Over-activation of excitatory neurons is signaled to the network through the elevation of BMP2, a growth factor well-known for its role as morphogen in embryonic development. BMP2 acts on parvalbumin-expressing (PV) interneurons through the transcription factor SMAD1, which controls an array of glutamatergic synapse proteins and components of peri-neuronal nets. PV interneuron-specific disruption of BMP2-SMAD1 signaling is accompanied by a loss of PV cell glutamatergic innervation, underdeveloped peri-neuronal nets, and decreased excitability. Ultimately, this impairment of PV interneuron functional recruitment disrupts cortical excitation – inhibition balance with mice exhibiting spontaneous epileptic seizures. Our findings suggest that developmental morphogen signaling is re-purposed to stabilize cortical networks in the adult mammalian brain.

## Introduction

Neuronal circuits in the neocortex underlie our ability to perceive our surroundings, integrate various forms of sensory information, and support cognitive functions. Cortical computation relies on assemblies of excitatory and inhibitory neuron types that are joined into canonical microcircuit motifs. The synaptic innervation and intrinsic properties of fast-spiking parvalbumin-expressing inhibitory interneurons (PV interneurons) have emerged as key parameters for controlling cortical circuit stability and plasticity ^1,2^. During development, sensory experience shapes synaptic innervation of PV interneurons in an afferent-specific manner and synaptic input to PV interneuron dendrites is a critical node for cortical dysfunction in disorders ^3–7^. In the adult brain, neuronal activity-dependent regulation of PV interneuron recruitment and excitability are fundamental for the maintenance of excitation – inhibition balance and have been implicated in gating cortical circuit plasticity during learning processes ^1,8–12^. However, the molecular mechanisms underlying these critical features, in particular trans-cellular signaling events that relay alterations in neuronal network activity and adjust PV interneuron function are poorly understood.

### Bone Morphogenetic Protein signaling is mobilized by neuronal network activity in adult neocortex

To identify candidate trans-cellular signals that are regulated by neuronal network activity in mature neocortical neurons, we examined secreted growth factors of the bone morphogenetic protein family (BMPs), which had been implicated in cell fate specification and neuronal growth during development ^13–23^. Amongst four bone morphogenetic proteins (BMP2,4,6,7) examined, *Bmp2* mRNA was significantly upregulated in glutamatergic neurons upon stimulation (3.5 +/- 0.5-fold, Extended Data Fig. 1a-d). A similar activity-dependent elevation of BMP2 was observed at the protein level in neurons derived from a *Bmp2* HA-tag knock-in mouse (*Bmp2^HA/HA^*, Extended Data Fig. 1e-g). As developmental morphogens, BMPs direct gene regulation in recipient cells through SMAD transcription factors (Fig. 1a) ^24–29^. Interestingly, the canonical BMP-target genes *Id1* and *Id3* were significantly upregulated in stimulated neocortical cultures, a process that was blocked by addition of the extracellular BMP-antagonist Noggin (Extended Data Fig. 1h, i). In the neocortex of adult mice, key BMP signaling components continue to be expressed, with the ligand BMP2 exhibiting highest mRNA levels in glutamatergic neurons (Extended Data Fig. 2a-c). To test whether BMP-target gene transcription is activated in response to elevated neuronal network activity in adult mice, we chemogenetically silenced upper layer PV interneurons in the barrel cortex (Fig. 1b). This local reduction of PV neuron-mediated inhibition results in increased neuronal network activity ^30,31^ accompanied by a 4- to 8-fold transcript increase for the activity-induced primary response genes *Fos* and *Bdnf* (Fig. 1c). Importantly, this chemogenetic stimulation also resulted in upregulation of the canonical SMAD1/5-dependent BMP target genes (*Id1*, *Id3*,) (Fig. 1c). We then mapped neuronal cell populations which exhibit BMP target gene activation in response to neuronal network activity with a novel temporally-controlled BMP-signaling reporter (BRX) (Fig. 1d). We combined BMP-response element sequences (4xBRE) from the *Id1* promoter ^32^ with the small molecule (LMI070)-gated miniX^on^ cassette ^33^ to drive a nucleus-targeted eGFP (Extended Data Fig. 3). Thus, the level of nuclear eGFP reports activation of BMP-signaling during a time window specified by LMI070 application (Extended Data Fig. 4). Chemogenetic stimulation resulted in a selective increase in the BRX-reporter activity in PV interneurons whereas mean reporter output in glutamatergic cells and non-PV interneurons was unchanged (Fig. 1f, g, but note that a sparse sub-population of NeuN^+^/Gad67^-^ glutamatergic neurons did show significant reporter signal). Genetic restriction of the BRX-reporter to PV-interneurons revealed a three-fold increase in BRX signal in response to chemogenetic stimulation (Fig. 1 h,i). In aggregate, these results demonstrate that increased cortical network activity mobilizes BMP2 and selectively activates the BMP-signaling pathway in PV interneurons in the adult mouse barrel cortex.

**Fig. 1.**
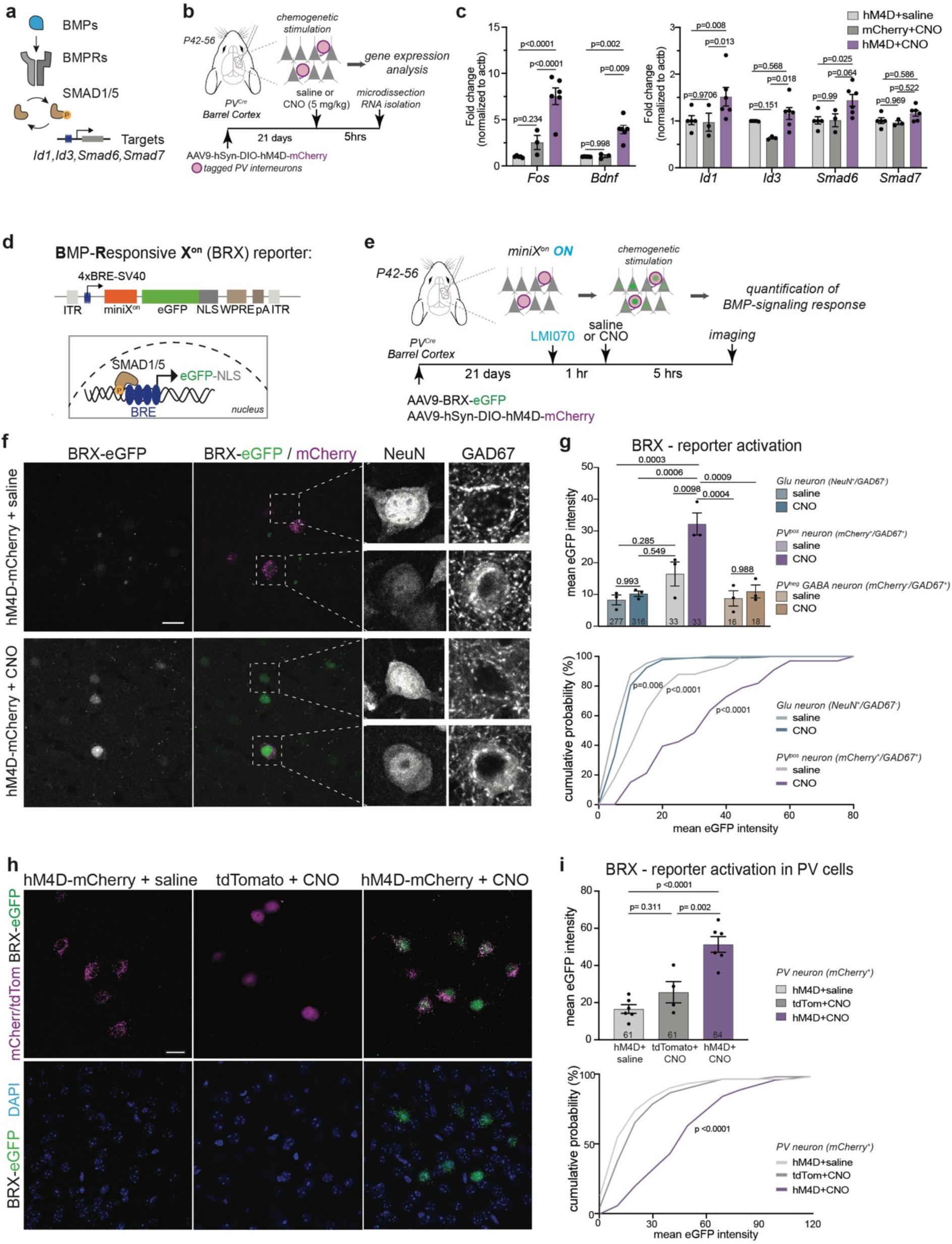
Neural activity elevation elicits BMP signaling in PV interneurons of the adult barrel cortex. (**a**) Illustration of BMP pathway components (adopted from ^35^). (**b**) Schematic representation of chemogenetic neuronal activity manipulation protocol in adult barrel cortex. **(c)** Expression of immediate early genes *cFos* and *Bdnf* and SMAD1/5 target genes *Id1*, *Id3*, *Smad6*, and *Smad7* in barrel cortex of chemogenetically stimulated and control mice (N=3-6 mice/group). Two-way ANOVA with Tukey’s post hoc test. **(d)** Schematic representation of the viral BRX reporter. Nucleus-targeted eGFP (NLS-eGFP) is expressed under control of regulatory elements from the *Id1* promoter (4xBRE) and the miniX^on^ splicing cassette. **(e)** Experimental paradigm. **(f)** Representative images of BRX reporter signal in barrel cortex layer 2/3 of *PV^Cre^* mice. Cre-dependent mCherry identifies PV cells, NeuN identifies neurons, somatic/perinuclear GAD67 signal identifies GABA neurons. **(g)** Quantification of BRX reporter readout in chemogenetically stimulated and control L2/3 neurons. Bar graph for mean ± SEM of nuclear eGFP intensity per mouse (N=3 mice/group, cell numbers indicated in columns, one-way Anova with Tukey’s multiple comparisons) and cumulative distribution of eGFP reporter intensity per PV interneuron (Kolmogorov-Smirnov test). **(h)** Representative images of cre-dependent BRX reporter selectively expressed in PV interneurons in barrel cortex layer 2/3. **(i)** Bar graph for mean ± SEM of nuclear eGFP intensity per mouse (N=4-6 mice/group, n=61-84 cells per condition, one-way Anova with Tukey’s multiple comparisons multiple comparisons) and cumulative distribution of eGFP reporter intensity per PV interneuron (Kolmogorov-Smirnov test). Scale bar in f and h is 20 μm.

### BMP-SMAD1 signaling controls transcriptional regulation of synaptic proteins

During development, the combinatorial action of various BMP ligands and receptors directs cell type-specific target gene regulation through SMAD transcription factors, but also SMAD-independent functions have been described ^13,16,19,21,34–39^. In neocortical neurons, BMP2 stimulation (20 ng/ml for 45 minutes) resulted in SMAD1/5 activation in both, glutamatergic and GABAergic neurons (Extended Data Fig. 5a-c). To uncover neuronal SMAD1 target genes, we performed chromatin immunoprecipitation sequencing (ChIP-seq) for SMAD1/5 from adult mouse neocortex and neocortical cultures (Fig. 2a). We identified 239 and 543 sites that were bound in mouse neocortex and cultured neocortical neurons, respectively (Fig. 2b and Supplementary Table 1). Notably, 77% of the binding sites in vivo were reproduced in the culture neuron preparations. To specifically map sites acutely regulated by BMP-SMAD1/5 signaling we stimulated cortical cultures by addition of recombinant BMP2. Upon stimulation, we identified another 353 BMP2-responsive SMAD1/5 binding sites. Importantly, the majority of BMP2-responsive peaks were associated with promoter elements. To explore whether SMAD1 triggers *de novo* activation of target genes or rather modifies transcriptional output of active genes, we mapped histone 3 acetylated at lysine 27 (H3K27ac) marks, a chromatin modification at active promoters and enhancers. By intersecting H3K27ac ChIP-seq signals with SMAD1/5 peaks (Fig. 2b-e), we found that the majority of BMP2-responsive elements contain significant H3K27ac marks that are slightly increased after stimulation. This suggests that many of these sites are already active without BMP2 stimulation. By comparison, constitutively bound regions exhibited lower H3K27ac signal (Fig. 2b, c). Sequence analysis identified enrichment of different motifs for SMAD1/5 DNA binding motif in the constitutive (tissue and neuronal culture) and BMP2-responsive gene regulatory elements, suggesting that DNA binding might involve different co-factors (Fig. 2d). The impact of BMP2-induced SMAD1/5 recruitment on transcriptional output was examined by RNA-sequencing. Differential gene expression analysis identified 30 and 147 up-regulated transcripts 1 and 6 hours after BMP2-stimulation, respectively (Extended Data Fig. 5d, Supplementary Table 2). 50% of the regulated genes 1 hour after BMP2-stimulation had direct SMAD1/5 binding at their promoters. These genes included known negative feedback loop genes of the BMP signaling pathway (*Id1*, *Id3* and *Smad7*). 25% of differentially regulated genes 6 hours after BMP2-stimulation had direct SMAD1/5 binding. (Extended Data Fig. 5d). Conditional knock-out of *Smad1* in post-mitotic neurons was sufficient to abolish upregulation of these genes in response to BMP2 signaling and reduced their expression in naïve (unstimulated) neurons (Fig. 2f and Extended Data Fig. 5e,f Supplementary Table 3). Direct transcriptional targets of BMP-SMAD1 signaling in neocortical neurons included an array of activity-regulated genes such as *Junb*, *Trib1* and *Pim3*, key components of the extracellular matrix (*Bcan, Gpc6*) and glutamatergic synapses (*Lrrc4*, *Grin3a*) (Fig.2e, Extended Data Fig. 4d). Moreover, neuronal *Smad1* ablation was accompanied by broad gene expression changes beyond de-regulation of direct SMAD1 target genes (Fig. 2g). Top GO terms enriched amongst the upregulated genes were “glutamatergic synapse” and transcription factors under the term “nucleus” (Fig. 2h). Furthermore, de-regulated genes include the majority of neuronal activity-regulated rapid primary (rPRG) and secondary (SRG) activity-response genes (Fig. 2i). Thus, SMAD1 is a key downstream mediator of BMP signaling in mature neurons and its neuronal loss of function results in a substantial upregulation of neuronal activity-response genes *in vitro*.

**Fig. 2.**
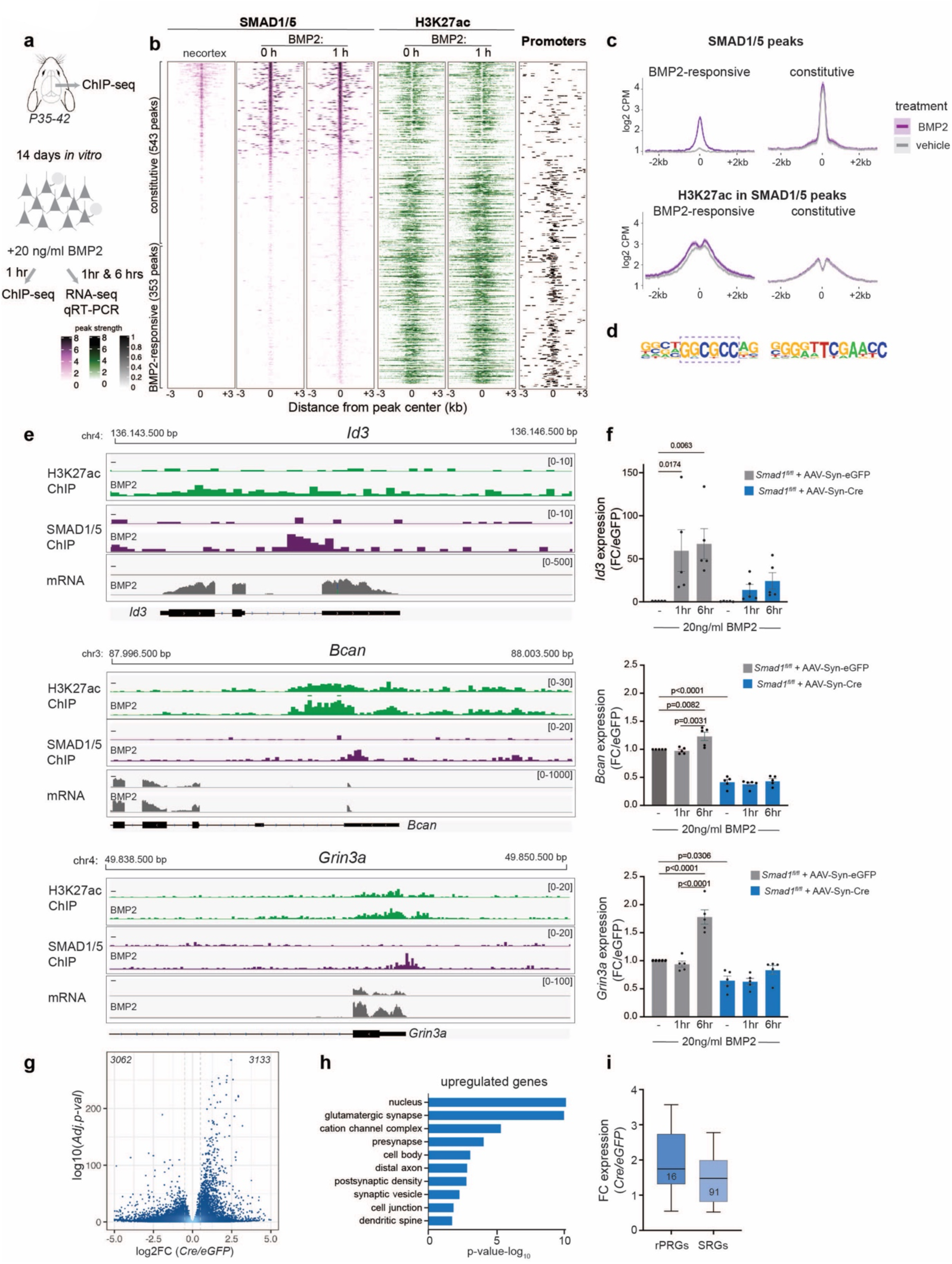
BMP2-SMAD1 signaling regulates synaptic components and is required for stable neuronal networks. **(a)** Schematic representation of ChIP-seq and RNA-seq experiments from mouse neocortex and neocortical cultures. **(b)** ChIP-seq analysis of neocortical tissue and naïve (0 hour) or growth factor-stimulated (1hour 20ng/ml BMP2) neocortical neuron cultures at DIV14. Heatmaps in purple display peak strength of SMAD1/5 binding, heatmaps in green show H3K27ac binding at SMAD1/5 peak regions. The right column (in black) displays position of promoter elements. Each binding site is represented as a single horizontal line centered at the SMAD1/5 peak summit, color intensity correlates with sequencing signal for the indicated factor. Peaks are ordered by decreasing Smad1/5 peak intensity. **(c)** Mean normalized ChIP-seq signal for SMAD1/5 and H3K27ac plotted for BMP2-responsive and constitutive SMAD1/5 binding sites. Gray lines indicate signal obtained from vehicle-treated cultures and purple lines signal obtained from BMP2-stimulated cultures. **(d)** Top enriched motifs detected for BMP2-responsive (left) and constitutive (right) SMAD1/5 peaks. **(e)** Examples of IGV genome browser ChIP-seq tracks displaying H3K27ac (green), SMAD1/5 (purple) and RNA-seq signal for SMAD1/5 targets *Id3*, *Bcan* and *Grin3a* in naïve (-) and BMP2-stimulated cultures. **(f)** qPCR analysis of mRNA expression of *Id3*, *Bcan* and *Grin3a* mRNAs in AAV-Syn-eGFP infected *versus* AAV-Syn-Cre infected *Smad1^fl/fl^* neocortical cultured neurons. Fold change (FC) relative to unstimulated cells is shown for 1 hour and 6 hours stimulation with 20ng/ml BMP2. **(g)** Volcano plot of differential gene expression in naïve *Smad1^fl/fl^* cortical cultures infected with AAV-Syn-iCre *versus* AAV-Syn-eGFP. Dashed lines indicate log_2_FC:0.4 and -log10Adj.-p-val: 2 chosen as thresholds for significant regulation. Number of significantly down- and up-regulated genes are indicated on the top. **(h)** Top ten enriched cellular component gene ontology terms for genes upregulated in conditional *Smad1* mutant cells (*Smad1^fl/fl^*infected with AAV-Syn-iCre) in unstimulated cortical cultures. **(i)** Expression levels of neuronal activity-regulated rapid Primary Response Genes (rPRGs) and Secondary Response Genes (SRGs) as defined in ^61^ in conditional *Smad1* mutant cells (*Smad1^fl/fl^* infected with AAV-Syn-iCre) compared to control AAV-Syn-eGFP infected cultures. The bar graphs show the means ± SEM (N=5 per condition, one-way ANOVA with Tukey’s multiple comparisons).

### Synaptic innervation and excitability of PV interneurons are controlled by SMAD1

In neocortical circuits excitation – inhibition balance is regulated by glutamatergic input synapses onto PV interneurons, and peri-neuronal nets (PNNs) surrounding these cells are modified in response to changes in neuronal network activity ^40,41^. To test whether pyramidal cell-derived BMP2 modifies PV interneuron innervation, we generated *Bmp2* conditional knock-out mice where *Bmp2* is selectively ablated in upper layer glutamatergic neurons (*Cux2^creERT2^:: Bmp2^fl/fl^*, referred to as *Bmp2^ΔCux2^* mice). We then adopted genetically encoded intrabodies (Fibronectin intrabodies generated by mRNA display, FingRs) to quantitatively map synaptic inputs to PV interneurons^42,43^ (Extended Data Fig. 6a-c and Supplementary Movie 1). A FingR-PSD95 probe was selectively expressed in PV interneurons in layer 2/3 of barrel cortex under control of a PV-cell-specific enhancer^44^ (Fig. 3a-d). Importantly, synapse density onto PV interneurons was reduced upon genetic ablation of *Bmp2* in upper layer pyramidal cells of *Bmp2^ΔCux2^* mice (Fig. 3e,f). We then generated PV interneuron-specific *Smad1* conditional knock-out mice to examine whether BMP2 acts through SMAD1. Postnatal ablation of *Smad1* (*PV^cre/+^::Smad1^fl/fl^*; referred to as *Smad1^ΔPV^* mice) did not alter PV cell density or distribution in the somatosensory cortex (Extended Data Fig. 7a-c). Using a cre recombinase-dependent form of the FingR-PSD95 probes (Fig. 4a-g), we observed a 40% reduction in morphological glutamatergic synapse density onto *Smad1^ΔPV^*interneurons (Fig. 4b, c). This was accompanied by a comparable reduction in mEPSC frequency but no change in mEPSC amplitude in acute slice recordings (Fig. 4d-f). The density of peri-somatic PV-PV synapses (identified by synaptotagmin-2 and a FingR-gephyrin probe ^42^) was also reduced (Fig. 4h, i). However, there was no significant change in mIPSC frequency or amplitude in PV cells of *Smad1^ΔPV^*mice, likely due to compensatory inhibition derived from other interneuron classes (Fig. 4j-l). Thus, SMAD1 is required for normal functional glutamatergic innervation of layer 2/3 PV interneurons, resulting in reduced glutamatergic input to these cells in *Smad1^ΔPV^* mice.

**Fig. 3.**
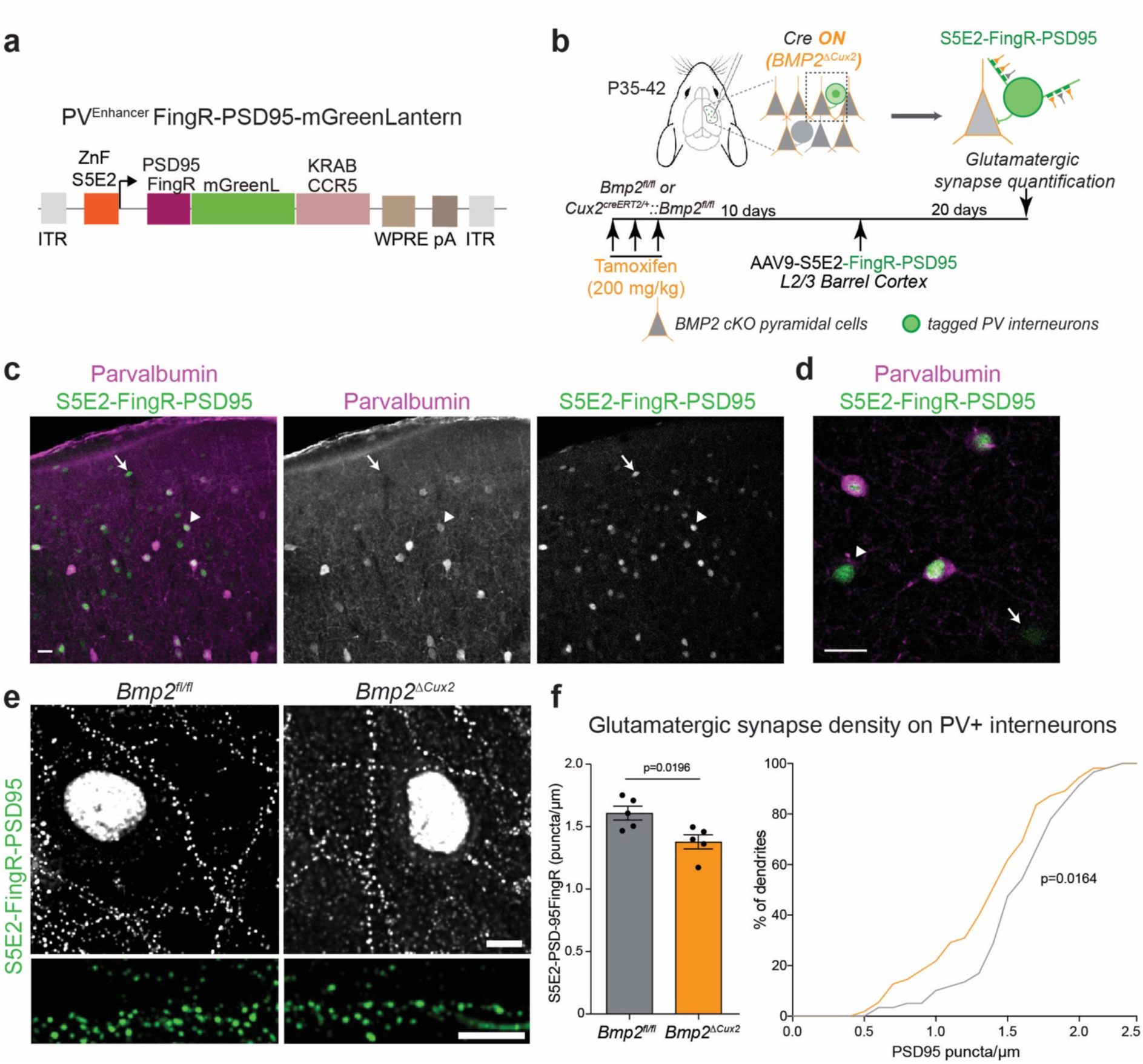
Pyramidal cell-derived BMP2 regulates PV interneuron innervation. **(a)** Schematic representation of the viral vector for expression of glutamatergic FingR-PSD95 probe. FingR expression is driven from S5E2 PV enhancer fused to a CCR5 zinc finger binding site (ZnF). FingR coding sequence is fused to mGreenLantern and a CCR5-KRAB transcriptional repressor for autoregulation of probe expression. Thus, excess probe accumulates in the nucleus. **(b)** Conditional deletion of BMP2 from upper layer neurons in *BMP2^ΔCux2^* mice is achieved by three tamoxifen applications spaced over one week. **(c,d)** Selectivity of FingR probe expression with S5E2 enhancer. A PV interneuron with low parvalbumin protein level is marked by an arrow head and a parvalbumin-negative cell is marked by an arrow. Scale bar is 20 μm **(e)** FingR-PSD95-marked synapses formed onto PV interneurons in control (*Bmp2^fl/fl^*) and *Bmp2^ΔCux2^*animals and corresponding dendritic stretches. **(f)** Quantification of glutamatergic synapse density on the dendrites of PV interneurons (identified by probe expression and parvalbumin immunostaining). Number of synapses were normalized to dendritic length (Mean and SEM from N=5 animals per genotype, n=7-17 dendrites per animal, unpaired t-test) and cumulative distribution of synapse density across all cells analyzed (n=55-60 cells, Kolmogorov-Smirnov test). Scale bar in e is 5 μm.

**Fig. 4.**
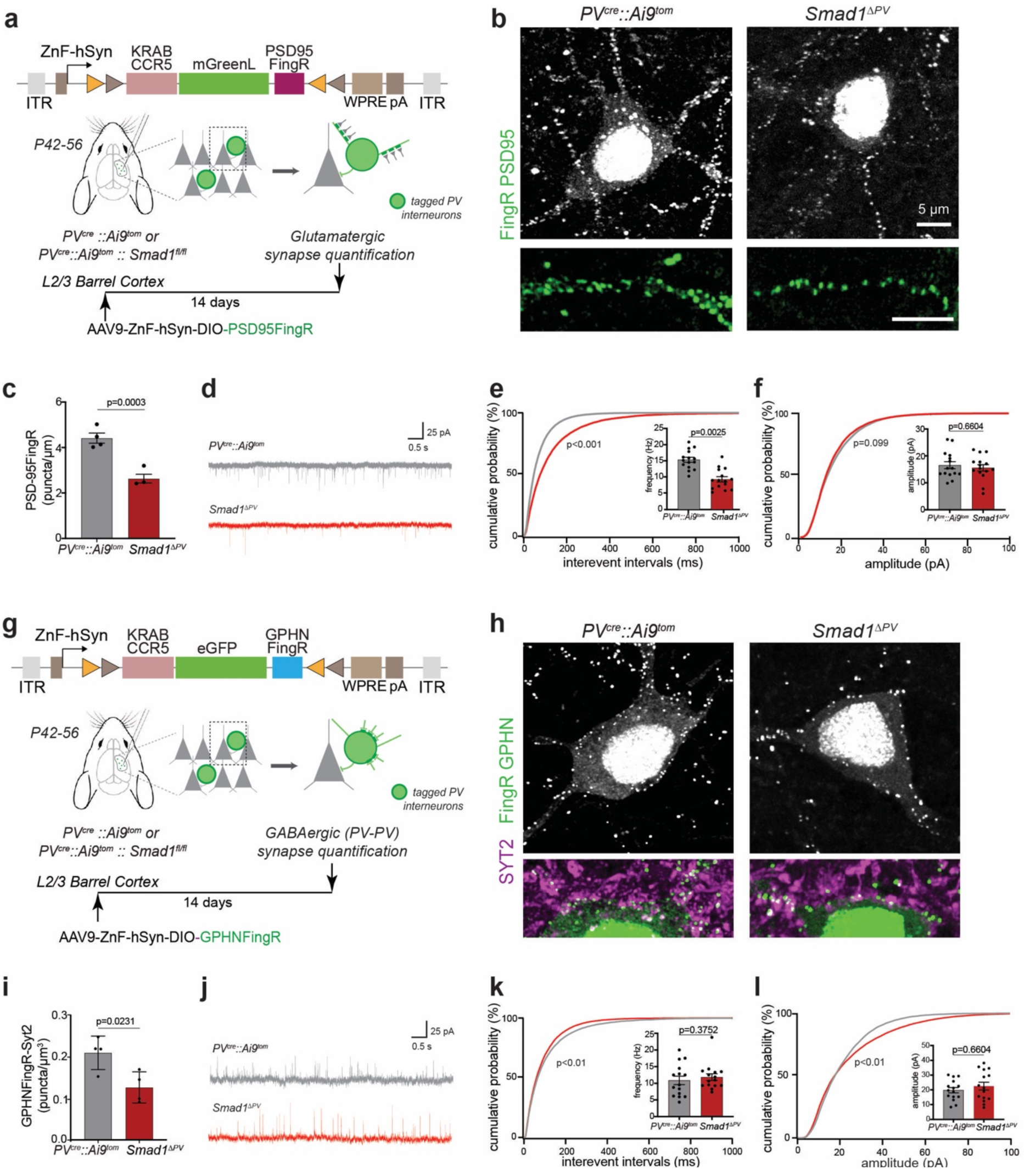
SMAD1 regulates glutamatergic innervation of PV interneurons. **(a)** Schematic representation of cre-recombinase-dependent intrabody probe. Intrabody expression is driven from human synapsin promoter (hSyn) fused to a CCR5 zinc finger binding site (ZnF). **(b)** FingR-PSD95-marked synapses formed onto control (*PV^cre^::Ai9^tom^*) and *Smad1* conditional knock-out (*Smad1^ΔPV^*) PV interneurons and corresponding dendritic stretches. **(c)** Quantification of glutamatergic synapse density on the dendrites of PV interneurons. Number of synapses was normalized to dendritic length (Mean and SEM from N=3-4 animals per genotype, n=10 dendrites per animal, unpaired t-test). Note that the vast majority of FingR-PSD95-marked structures co-localize with the presynaptic marker vGluT1 (see Extended Data Figure 6A). **(d)** Representative traces of mEPSC recordings from control (gray) and *Smad1^ΔPV^* (red) PV interneurons in acute slice preparations from adult mice. **(e)** Frequency distribution of interevent intervals (Kolmogorov-Smirnov test) and mean mEPSC frequency (mean ± SEM for n=15 cells/genotype, from N=3-4 mice. Kolmogorov-Smirnov test). **(f)** Frequency distribution of mEPSC amplitudes (Kolmogorov-Smirnov test) and mean mEPSC amplitude (mean ± SEM for n=15 cells/genotype, from N=3-4 mice. Kolmogorov-Smirnov test). **(g)** Schematic representation of AAV-driven, cre-recombinase-dependent intrabody probes for GABAergic synapses (GPHN-FingR), fused to eGFP and a CCR5-KRAB transcriptional repressor for autoregulation of probe expression. Thus, excess probe accumulates in the nucleus. **(h)** Synapses formed onto control (*PV^cre^::Ai9^tom^*) and *Smad1* conditional knock-out (*Smad1^ΔPV^*) PV interneurons. **(i)** Quantification of PV-PV GABAergic synapse density on PV interneuron somata. Number of GPHNFingR-eGFP / Synaptotagmin2 (SYT2) – containing structures were normalized to soma volume (mean and SEM from N=3-4 animals per genotype, n=78 cells, unpaired t-test). **(j)** Representative traces of mIPSCs recorded from control (in gray) and *Smad1^ΔPV^*(red) PV interneurons in acute slice preparations. **(k)** Frequency distribution of interevent intervals (Kolmogorov-Smirnov test) and mean mIPSC frequency (mean ± SEM for n=15 cells/genotype, from N=3-4 mice. Kolmogorov-Smirnov test). **(l)** Frequency distribution of mIPSC amplitudes (Kolmogorov-Smirnov test) and mean mIPSC amplitude (mean ± SEM for n=15 cells/genotype, from N=3-4 mice. Kolmogorov-Smirnov test). Scale bar in b and h is 5 μm.

Neuronal activity-induced regulation in PV interneurons modifies the elaboration of PNNs and parvalbumin expression ^2,30,40,41,45,46^, and our ChIP-Seq analysis identified the PNN component brevican (*Bcan*) as one of the direct SMAD1 targets. In *Smad1^ΔPV^* mice, the elaboration of PNNs around PV interneurons and parvalbumin protein expression were significantly reduced (Fig. 5a-c, and Extended Data Fig. 7d-g). Conversely, PV cell-specific activation of the BMP-signaling pathway by expression of a constitutively active BMP-receptor was sufficient to elevate parvalbumin levels (Extended Data Fig. 8). Through organizing PNNs, brevican has been implicated in regulating plasticity and excitability of PV interneurons ^41^. Interestingly, the firing rate of SMAD1-deficient PV interneurons in response to current injections was significantly reduced in the barrel cortex of adult mice (Fig. 5d-f and Extended Data Fig. 9a, note that firing rate as well as mEPSC frequency was unchanged in young animals, Extended Data Fig. 9b-e). This reduced firing frequency most likely is explained by a reduction in input resistance (IR) in the *Smad1^ΔPV^* cells (Extended Data Fig. 9a). Thus, in the absence of BMP-SMAD1 signaling PV interneurons not only receive less glutamatergic drive but are also less excitable. These cellular alterations resulted in a severe overall disruption of cortical excitation – inhibition balance. As compared to control littermates, *Smad1^ΔPV^* mice exhibited hyperactivity in open field tests and frequently exhibited spontaneous seizures when introduced into novel environments (Fig. 5g, h). Video-coupled EEG recordings with electrodes over the barrel cortex (Supplementary Movie 2) revealed marked high amplitude activity bursts at the time of seizure followed by a refractory period (Fig. 5i). Overall, our results demonstrate that elevated network activity in the somatosensory cortex of adult mice triggers the upregulation of BMP2 in glutamatergic neurons which balances excitation by controlling synaptic innervation and function of PV interneurons through the transcription factor SMAD1.

**Fig. 5.**
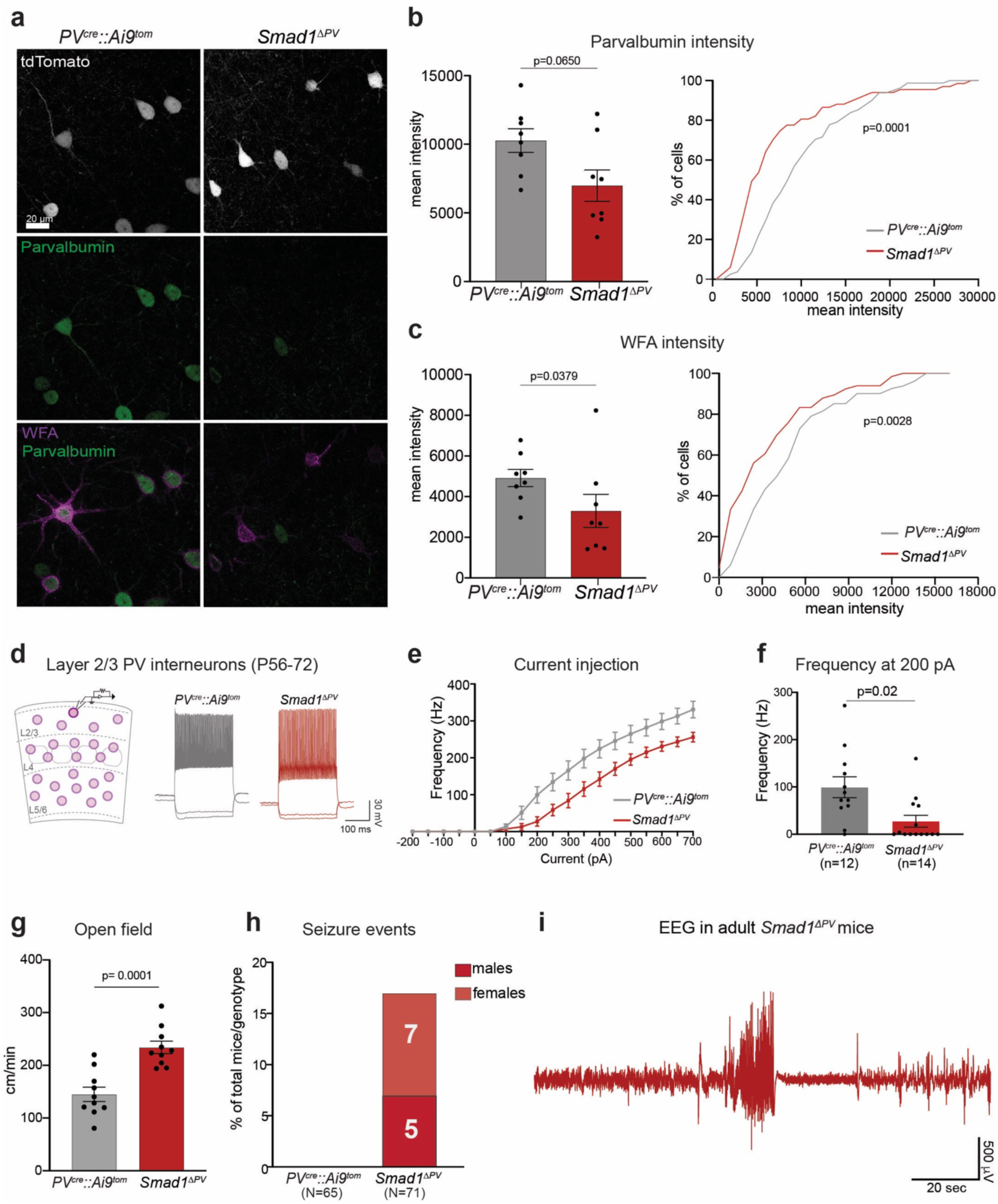
Loss of SMAD1 in PV interneurons results in disruption of E/I balance in the adult mice. **(a)** Parvalbumin immunoreactivity and Wisteria floribunda agglutinin (WFA)-binding to the PNNs in adult control (*PV^cre^::Ai9^tom^*) and *Smad1* conditional knock-out (*Smad1^ΔPV^*) mice. **(b)** Quantification of parvalbumin immunoreactivity per cell in *PV^cre^::Ai9^tom^*(gray) and *Smad1^ΔPV^* (red) mice. Bar graphs with mean intensity per mouse (N=8/genotype) and cumulative distribution of mean intensity per cell (n=81 cells for *PV^cre^::Ai9^tom^*, n=67 cells for *Smad1^ΔPV^* mice). Kolmogorov-Smirnov test for bar graph and cumulative distribution. **(c)** WFA staining intensity plotted as in (b). **(d)** Experimental strategy and example traces from current-clamp recordings of control (in gray) and *Smad1^ΔPV^* (red) PV interneurons in acute slice preparations. **(e)** Comparison of firing frequencies of layer 2/3 PV interneurons at given currents and **(f)** Mean firing frequency in response to 200 pA current injection in cells from *PV^cre^::Ai9^tom^* (gray) and *Smad1^ΔPV^* (red) mice (N=4 mice, n=12 cells for *PV^cre^::Ai9^tom^*and N=4, n=14 cells for *Smad1^ΔPV^*, Kolmogorov-Smirnov test). **(g)** Quantification of the velocity in open field from adult *PV^cre^::Ai9^tom^*(gray) and *Smad1^ΔPV^* (red) mice (N=10 mice/genotype, unpaired t-test). **(h)** Number of *PV^cre^::Ai9^tom^* control (0 out of 65 mice) and male and female *Smad1^ΔPV^* (red) mice (12 out of 71 mice) displaying spontaneous seizures during cage changes. Note that observation of spontaneous seizures was an exclusion criterion for morphological, electrophysiological or molecular analyses performed in this study. **(i)** Representative 2.5 minutes EEG trace obtained from a *Smad1^ΔPV^* mouse. All bar graphs show the means ± SEM.

## Discussion

Despite being exposed to a wide range of sensory stimulus intensities, cortical circuits exhibit remarkably stable activity patterns that enable optimal information coding by the network. This network stability is achieved by homeostatic adaptations that modify the excitability of individual neurons, scale the strength of synapses, as well as microcircuit-wide modifications of excitatory and inhibitory synapse density ^10,12,47–50^. These multiple adaptations occur at various time-scales, from near instantaneous adjustments of excitation and inhibition during sensory processing ^51^, to slower modifications of synaptic connectivity upon longer-term shifts in circuit activation as they occur during sensory deprivation but also in disease states ^12,52–56^. Thus, both rapid cell intrinsic, as well as long-lasting trans-cellular signaling processes have evolved to ensure cortical network function and stability.

We here demonstrate that elevated neuronal network activity in the somatosensory cortex of adult mice triggers BMP2 upregulation in pyramidal cells and BMP-target gene expression in PV interneurons. The transcription factor SMAD1, directly binds to and regulates promoters of an array of glutamatergic synapse proteins and components of the perineuronal nets, such as brevican. Thus, BMP2-SMAD signaling provides a trans-neuronal signal to adjust functional PV interneuron recruitment and excitability that ultimately serve to maintain excitation – inhibition balance and stabilize cortical network function in adult neocortex. Importantly, the SMAD1 loss-of-function phenotypes only emerge with age, as PV interneuron excitability and synaptic innervation are normal in adolescent (P26-P30) mice. In developing auditory cortex, genetic deletion of type I BMP-receptors from PV interneurons is associated with impaired synaptic plasticity at PV interneuron output synapses onto principal neurons of layer 4 whereas basal GABAergic transmission was unchanged ^57^. This suggests a selective role for BMP2-SMAD1 signaling in controlling glutamatergic input connectivity to PV interneurons.

Importantly, transcriptional regulation through BMP2-SMAD1 signaling significantly differs from the action of activity-induced immediate early genes. As secreted growth factor, BMP2 derived from glutamatergic neurons relays elevated network activity to PV interneurons through the activation of an array of SMAD1 target genes. Rather than ON/OFF responses, the majority of direct SMAD1 targets exhibit active enhancer and promoter elements and are already expressed under basal conditions. However, SMAD1 activation results in an elevation of transcriptional output, indicating a graded gene expression response to BMP2.

In early development, BMP growth factors act as morphogens that carry positional information and differentially instruct cell fates ^26,27,29^. The combinatorial complexity arising from the substantial number of BMP ligands and receptors has the power to encode computations for finely tuned cell-type-specific responses ^35,58^. Our work suggests that the spatiotemporal coding power, robustness, and flexibility which evolved for developmental patterning is harnessed for balancing plasticity and stability of neuronal circuits in the adult mammalian brain. Interestingly, additional BMP ligands besides BMP2 are selectively expressed in neocortical cell types (Extended Data Fig. 2b). Moreover, an array of type I and type II BMP receptors are detected across neocortical cell populations. Thus, BMP-SMAD1 signaling might control additional aspects of neuronal cell-cell communication.

Disruptions in excitation – inhibition balance and homeostatic adaptations have been implicated in neurodevelopmental disorders as there is reduced GABAergic signaling and a propensity to develop epilepsy in individuals with autism ^54–56,59^. Considering that BMP-signaling pathways can be targeted with peptide mimetics ^60^ they may provide an entry point for therapeutic interventions in neurodevelopmental disorders characterized by disruptions in PV interneuron innervation, excitation – inhibition balance, and seizures.

## Acknowledgements

We thank Gustavo Aguilar, Shirley Dixit, Wuzhou Yang, Özgür Genç, Zayna Chaker, Josef Bischofberger, Filippo Rijli, Ralf Schneggenburger and Kelly Tan for support, advice and valuable comments on the manuscript. Caroline Bornmann, Sabrina Innocenti, Enrique Perez-Garci provided excellent technical assistance. We are grateful to the Biozentrum Imaging Core Facility, the Centre for Transgenic Models, and the Quantitative Genomics facility Basel for expert support. This work was financially supported by a Fellowship of Excellence from Biozentrum, University of Basel to Z.O., an EMBO Long-term Fellowship to M.P (ALTF 672-2022), an EMBO Long-term Fellowship to R.O. (ALTF 378-2020), and Grants from the Swiss National Science Foundation to P.S. (grant # 179432, # 154455, and # 209273). The Scheiffele Laboratory is an associated member of the NCCR RNA & Disease.

## Author contributions

All authors contributed to the design and analysis of experiments. Z.O. and N.S. conducted genetic and in vivo manipulations, Z.O. and M.P. conducted electrophysiological recordings, R.O. and M.S. contributed to analysis of ChIP and RNA Seq data, V.B. conducted EEG recordings, Z.O., D.S., N.S. and K.K. performed molecular biology procedures. Z.O. and P.S. wrote the manuscript with feedback and editing provided by all authors.

## Materials & correspondence

ChIP-seq and gene expression data will be deposited at GEO. All renewable reagents will be distributed by the corresponding author or deposited in public repositories for distribution. The regulatory elements of the X^on^ system and cDNA sequences encoding FingRs will be distributed in accordance with respective MTAs under which they were originally obtained by the authors.

## Methods

### Animals

All procedures involving animals were approved by and performed in accordance with the guidelines of the Kantonales Veterinäramt Basel-Stadt. All experiments were performed in mice in C57Bl/6J background, except for some of the experiments performed in cultured wild-type neurons which used RjOrl:SWISS mice. Both males and females were used at similar numbers for the experiments. Animals were randomly assigned to treatment groups. Animals that exhibited a spontaneous seizure were excluded from molecular, anatomical, or slice physiology analysis.

*Smad1^floxed^ mice* ^62^*, Pvalb-cre* mice ^63^ and *Ai9* mice ^64^ were obtained from Jackson Laboratories (Jax stock no: 008366, 017320, and 007909 respectively). Cux2-CreERT2 mice ^65^ were obtained from the Mutant Mouse Resource and Research Center (MMRRC). *Bmp2*-2xHA mice were generated using a Crispr-Cas9 strategy ^66^ inserting a double HA tag at the N-terminus of the mature BMP2 protein, between amino acids S292 and S293. Guide RNAs (gRNAs) employed were 5’-GTCTCTTGCAGCTGGACTTG-3’ and 5’-CAAAGGGTGTCTCTTGCAGC-3’, together with a 200 bp single-stranded DNA ultramer: 5’-GACTTTTGGACATGATGGAAAAGGACATCCGCTCCACAAACGAGAAAAGCGTCAAGCC AAACACAAACAGCGGAAGCGCCTCAAGTCCGCTAGC**TACCCATACGATGTTCCAGATT ACGCT**GGCTATCCCTATGACGTCCCGGACTATGCAGCTAGCAGCTGCAAGAGACACCC TTTGTATGTGGACTTCAGTGATGTGG-3’ (sequencing encoding the HA tags highlighted in bold).

### Surgery and drug treatments

Injections of recombinant AAVs were performed into the barrel cortex of 42-49 days old male and female mice performed under isoflurane anesthesia (Baxter AG, Vienna, Austria). Mice were placed in a stereotactic frame (Kopf, Germany) and a small incision (0.5 – 1 cm) was made over the barrel cortex at the following coordinates targeting two sites: ML ±3.0 mm and ±3.4 mm, at AP 0.6 mm and AP -1.6 mm, DV –1.5 mm from Bregma to target layer 2/3 and layer 4). For injections of FingR intrabodies, two injection sites restricted to layer 2/3 were used: ML ±3.0 mm and ±3.4 mm at AP –1.0 mm, DV –0.96 mm from Bregma. Recombinant AAVs (titer: 10^12^-10^13^) were injected via a glass capillary with outer diameter of 1 mm and inner diameter of 0.25 mm (Hilgenberg) for a total volume of 100 nL per injection site. The wound was closed with sutures (Braun, # C0766194).

LMI070 (25 mg/kg, MedChemExpress, HY-19620, suspended in 20% cyclodextrin and 10% DMSO to 5 mg/ml concentration) was administrated by oral gavage. Clozapine N-oxide (5 mg/kg CNO, Sigma Aldrich, C0832) and doxycycline (50 mg/kg, FisherScientific, BP26531, suspended in 0.9% NaCl to 5 mg/ml concentration) were administered by intraperitoneal injection.

### Antibodies and probes

Monoclonal mouse-anti-Synaptotagmin 2 (Zebrafish International Resource Center, # ZNP-1), rabbit anti-Smad1 (Cell Signaling #9743), H3K27ac (Abcam #4729), rabbit anti-SMAD5 (Cell Signaling, #12534), anti-phosphoSMAD1/5/9 (Cell signaling #13820), mouse anti-BMPR2 (BD Pharmingen, #612292), rabbit anti-Calnexin (StressGen, SPA-865), mouse anti-MAP2 (Synaptic systems, 188011), mouse anti-CAMKII alpha (ThermoFisher, MA1-048), rat anti-GAPDH (Biolegend, 607902), rabbit anti-NeuN (Abcam, ab177487), mouse anti-GAD67 (Millipore #MAB5406), rabbit-anti-vGlut1 (Synaptic Systems #135303), rabbit IgG (Cell Signaling Technology #2729 as control IgG for ChIP) biotinylated WFA (Vector laboratories #B-1355-2), rabbit-anti-HA (Cell Signaling #3724), mouse anti-eGFP antibody (Santa Cruz, sc-9996), goat anti-Parvalbumin antibody (Swant #PVG213). Secondary antibodies were: HRP goat anti-rabbit (Jackson #111-035-003), HRP goat anti-rat (Jackson #112-035-143), HRP goat anti-mouse (Jackson #115-035-149), Alexa405 goat anti-rabbit (Thermo Scientific #A-31556), Alexa488 donkey anti rabbit (ThermoFisher #R37118), Alexa647 donkey anti-mouse (Jackson #715-605-151), Alexa647 Streptavidin (Thermo Scientific, S32357), Cy2 Streptavidin (Jackson #016-220-084), Cy3 donkey anti-mouse (Jackson #715-165-151), Cy3 donkey anti-rabbit (Jackson #711-165-152), Cy3 donkey anti-guineapig (Jackson #706-165-148), Cy5 donkey anti-goat (Jackson #705-175-147), Cy5 donkey anti-rabbit (Jackson #711-175-152) and Cy5 donkey anti-mouse (Jackson #715-175-511). DAPI dye was used for nuclear staining (TOCRIS bio techne^®^, 5748).

### Immunohistochemistry and image acquisition

Mice were deeply anesthetized with ketamine/xylazine (100/10 mg/kg i.p.) and transcardially perfused with fixative (4% paraformaldehyde in 0.1 M phosphate buffer, pH 7.4). For synapse quantifications with FingR probes fixative also contained 15% picric acid. After perfusion, brains were post-fixed overnight in fixative at 4°C, washed 3 times with 100 mM phosphate buffer (PB).

For quantifications of parvalbumin and WFA expression and BRX reporter analyses, coronal brain slices were cut at 40 µm with a Vibratome (VT1000S, Leica). For FingR-PSD95 analysis with the cre-dependent reporter, coronal brain slices were cut at 30 µm with a Cryostat (Microm HM560, Thermo Scientific). Brain sections were incubated for 30 minutes in blocking solution (0.3% Triton X-100 and 3% Bovine Serum Albumin in PBS). Sections were incubated with primary antibodies in blocking solution overnight at 4°C, washed three times (10 minutes each) with 0.05% Triton X-100 in PBS, followed by incubation for 1.5 hours at room temperature with secondary antibodies in blocking solution. Sections were washed three times with PBS and DAPI dye (1.0 µg/ml) co-applied during the wash. Sections were mounted using Microscope cover glasses 24×60mm (Marienfeld Superior™ 0101242) on Menzel-Gläser microscope slides SUPERFROST^®^ PLUS (Thermo Scientific, J1800AMNZ) with ProLongTM Diamond Antifade Mountant (InvitrogenTM, P36970).

For S5E2 PV enhancer FingR-PSD95 quantifications, coronal brain slices were cut at 120 µm on a Vibratome (VT1000S, Leica) and cleared with CUBIC-L solution (10% w/v N-Butyldiethanolamine, 10% w/v Triton-X-100) for 3 hours at 37 °C with gentle shaking^67^. Sections were stained with goat anti parvalbumin antibodies and mounted with Ce3D™ Tissue Clearing Solution (Biolegend, 427704).

For parvalbumin and WFA analysis, images were acquired on an inverted LSM700 confocal microscope (Zeiss) using 20x/0.45 and 40X/1.30 Apochromat objectives. For cell density quantifications of PV interneurons, tilescan images from Barrel Cortex were acquired. For synapse quantifications, images were acquired with a PlanApo 63x/1.4 oil immersion objective.

For primary neocortical neurons in culture, fixation was with 4% PFA in 1X PBS for 15min. followed by ice-cold methanol (10min at -20°C). Cells were blocked (5% donkey serum, 0.3% Triton X-100 in PBS) for 1hr at room temperature and primary antibody incubation was performed overnight at 4°C in a humidified chamber. Secondary antibody incubation was 1hr at room temperature. Imaging was performed on a widefield microscope (Deltavision, Applied Precision) with a 60X objective (NA 1.42, oil).

### Image analysis

Mean intensity analyses for parvalbumin and WFA stainings were performed in ImageJ with a custom-made script in Python. Briefly, H-Watershed was applied to segment PV interneurons based on tdTomato signal on the soma. To detect WFA signal, the soma was eroded and dilated in all optical sections. After applying thresholding, parvalbumin and WFA mean intensity values were automatically calculated and displayed as arbitrary units. Integrity analysis of perineuronal nets was done from PV interneurons with positive WFA signal (>2000 arbitrary unit). Images were post-processed by conservative deconvolution with the Huygens Deconvolution software with the classic maximum likelihood estimation deconvolution algorithm. Quantitative analysis of number of peaks and the distance between the peaks were performed by using plot profile function in Image J as described ^68^.

For BRX-reporter experiments, cell identity and reporter intensity were quantified with ImageJ. A region of interest was drawn around the nuclei (marked by DAPI) and mean intensity was measured for nuclear GFP signal and normalized to background fluorescence in the same image. Cells were identified based on immunostaining for markers (mCherry genetically restricted to PV interneurons), NeuN (marking neurons with high intensity in pyramidal cells), GAD67 (marking all GABAergic cells).

For synapse quantification, images were post-processed by conservative deconvolution with the Huygens Deconvolution software with the classic maximum likelihood estimation deconvolution algorithm. Quantitative analysis was performed using Imaris 9.8 by application of spots and surface detection tool.

All data collection and image analysis were done blinded to the genotype or treatment of the animal. Statistical analyses were done with Prism 9 (GraphPad software). Images were assembled using ImageJ and Adobe Illustrator software.

### ChIP-seq analysis

For ChIP-seq analysis with cultured neurons, 24×10^6^ cells (DIV14) were cross-linked with 1% formaldehyde for 10 min at room temperature. Crosslinking was stopped by the addition of glycine solution (Cell Signaling Technology, #7005) for 5 min at room temperature. Cells were scraped, pelleted, and lysed for 10 min on ice in 100 mM HEPES-NaOH pH 7.5, 280 mM NaCl, 2 mM EDTA, 2 mM EGTA, 0.5% Triton X-100, 1% NP-40, 20% glycerol. Nuclei were pelleted by centrifugation, washed in 10 mM Tris-HCl pH 8.0, 200 mM NaCl, and suspended in 10 mM Tris-HCl pH 8.0, 100 mM NaCl, 1 mM EDTA, 0.5 mM EGTA, 0.1% Na-Deoxycholate, 0.5% N-Lauroylsarcosine. Chromatin was sheared using a Covaris Sonicator for 20 minutes in sonication buffer (SimpleChIP Plus Sonication Kit, Cell Signaling Technology, #57976) to obtain fragments in the range of 200-500 bp. After sonication, sheared chromatin was centrifuged at 16,000 x g for 20 minutes at 4 °C and dissolved in 1x ChIP buffer (Cell Signaling Technology, #57976). 2% input was taken and chromatin was incubated with antibodies overnight at 4°C. Incubation with Protein G magnetic beads, decrosslinking and elution were performed as described in the SimpleChIP Plus Sonication Kit.

Libraries were generated using the KAPA Hyper Prep (Roche #KK8504) according to the manufacturer’s instructions and PCR amplified. Library quality was assessed using the High Sensitivity NGS Fragment Analysis Kit (Advanced Analytical #DNF-474) on the Fragment Analyzer (Advanced Analytical). Libraries were sequenced Paired-End 41 bases on NextSeq 500 (Illumina) using using 2 NextSeq 500 High Output Kit 75-cycles (Illumina, Cat# FC-404-1005) loaded at 2.5pM and including 1% PhiX. Primary data analysis was performed with the Illumina RTA version 2.4.11 and Basecalling Version bcl2fastq-2.20.0.422. Two Nextseq runs were performed to compile enough reads (on average per sample in total: 50±2 millions pass-filter reads).

ChIP-seq analysis from P35-42 mouse cortex was performed using SimpleChIP Enzymatic Chromatin IP Kit (Cell Signaling Technology, #9003) following manufacturer’s instructions with slight modifications. Briefly, neocortices from both hemispheres were crosslinked in 1.5% formaldehyde for 20 minutes at room temperature. Crosslinking was stopped by the addition of glycine solution for 5 minutes at room temperature. Tissue was pelleted, washed and disaggregated by using a Dounce homogenizer in 1x PBS containing protease inhibitor cocktail. Nuclei were pelleted by centrifugation and chromatin was digested by using micrococcal nuclease for 20 minutes at 37°C by frequent mixing to obtain fragments in the range of 150-900 bp. Nuclei were pelleted, resuspended in 1x ChIP buffer, sonicated with Bioruptor Pico (Diagenode #B01060010) to release sheared chromatin and was centrifuged at 9,400 x g for 10 minutes at 4°C. 2% input was taken and chromatin was incubated with primary antibodies overnight at 4°C. After subsequent incubation with 30 μl Protein G magnetic beads for 2 hours at 4°C, beads were washed three times with low salt, one time with high salt, one time with NP-40 buffer (8 mM Tris-HCl pH 8.0, 2 mM LiCl, 0.8 mM EDTA, 0.4% NP-40, and 0.4% sodium-deoxycholate) and one time with TE buffer (10 mM Tris-HCl, pH 8.0, 1 mM EDTA) at 4°C. Decrosslinking and elution were performed as described in the Enzymatic Chromatin IP Kit. Libraries were generated using the NEBNext® Ultra™ II DNA Library Prep Kit for Illumina (New England Biolabs, #E7645L) according to the manufacturer’s instructions and PCR amplified. Library quality was assessed using the High Sensitivity NGS Fragment Analysis Kit (Advanced Analytical, #DNF-474) on the Fragment Analyzer (Advanced Analytical, Ames, IA, USA) and cleaned up by using 1.0x Vol SPRI beads (Beckman Coulter). Libraries were sequenced Paired-End 41 bases on NextSeq 500 (Illumina) using 2 NextSeq 500 High Output Kit 75-cycles (Illumina, Cat# FC-404-1005). Two Nextseq runs were performed to compile enough reads (19-32 million pass-filter reads).

### RNA library preparation and sequencing

Libraries of BMP2-stimulated naïve cortical cultures were prepared from 200 ng total RNA by using the TruSeq Stranded mRNA Library Kit (Cat# 20020595, Illumina, San Diego, CA, USA) and the TruSeq RNA UD Indexes (Cat# 20022371, Illumina, San Diego, CA, USA). 15 cycles of PCR were performed.

Quality check was performed by using the Standard Sensitivity NGS Fragment Analysis Kit (Cat# DNF-473, Advanced Analytical) on the Fragment Analyzer (Advanced Analytical, Ames, IA, USA) and quantified (average concentration was 213±15 nmol/L and average library size was 357±8 base pairs) in order to prepare a pool of libraries with equal molarity. The pool was quantified by Fluorometry using using the QuantiFluor ONE dsDNA System (Cat# E4871, Promega, Madison, WI, USA) on Quantus instrument (Promega). Libraries were sequenced Single-reads 76 bases (in addition: 8 bases for index 1 and 8 bases for index 2) on NextSeq 500 (Illumina) using the NextSeq 500 High Output Kit 75-cycles (Illumina, Cat# FC-404-1005). Flow lanes were loaded at 1.4pM of pool and including 1% PhiX. Primary data analysis was performed with the Illumina RTA version 2.4.11 and Basecalling Version bcl2fastq-2.20.0.422. The Nextseq runs were performed to compile on average per sample: 56±3 millions pass-filter reads (illumina PF reads).

For the libraries from control and Smad1 mutant primary cortical cultures (four biological replicates), 100 ng total RNA was used and library preparation and quality check were performed as described above. Quantification yielded average concentration as 213±15 nmol/L and average library size as 357±8 base pairs. Libraries were sequenced Paired-End 51 bases (in addition: 8 bases for index 1 and 8 bases for index 2) setup using the NovaSeq 6000 instrument (Illumina). SP Flow-Cell was loaded at a final concentration in Flow-Lane loaded of 400pM and including 1% PhiX. Primary data analysis was performed as described above and 43±5 millions per sample (on average) pass-filter reads were collected on 1 SP Flow-Cell.

### ChIP- and RNA-seq data analysis

ChIP-seq reads were aligned to the December 2011 (mm10) mouse genome assembly from UCSC ^69^. Alignments were performed in R using the qAlign function from the QuasR package1 (v. 1.14.0) with default settings^70^. This calls the Bowtie aligner with the parameters “ -m 1 –best –strata”, which reports only reads that map to a unique position in the genome2. The reference genome package (“BSgenome.Mmusculus.UCSC.mm10”) was downloaded from Bioconductor (https://www.bioconductor.org). BigWig files were created using qExportWig from the QuasR package with the bin size set to 50. Peaks were called for each ChIP replicate against a matched input using the MACS2 callpeak function with the default options. Peaks were then annotated to the closest gene and to a genomic feature (promoter, 3’ UTR, exon, intron, 5’ UTR or distal intergenic) using the ChIPseeker R package. The promoter region was defined as -3 kb to + 3kb around the annotated TSS. Transcripts were extracted from the TxDb.Mmusculus.UCSC.mm10.ensGene annotation R package. All analyses in R were run in RStudio v. 1.1.447 running R v. 3.5.1. Enrichment of Bmp2-induced peaks over constitutive peaks was analyzed by using default settings in voom/limma analysis software packages. Motif enrichment analysis for BMP2-responsive peaks and constitutive peaks was performed separately by screening for the enrichment of known motifs with default settings of HOMER^71^. Output motif results with the lowest p-value and highest enrichment in targets compared to the background sequences were displayed for each peak set.

RNA-seq reads were aligned to mm10 using STAR and visualized in IGV genome browser to determine strand protocol. By using QuasR’s qQCReport, read quality scores, GC content, sequence length, adapter content, library complexity and mapping rate were checked and QC report was generated. Reads with quality scores less than 30, and mapping rates below 65 and having contaminations from noncoding RNAs were not considered for further analysis. If passed QC, QuasR’s qCount function was used to count reads mapping to annotated exons (from Ensembl genome annotations). Each read was counted once based on its start (if reads are on the plus strand) or end (if reads are on the minus strand) position. For each gene, counts were summed for all annotated exons, without double-counting exons present in multiple transcript isoforms (“exon-union model”). Correlations between replicates and batch structure were checked by plotting correlation heatmap, PCA plot of samples, scatter plots of normalized read counts. EdgeR package from R was used to build a model and test for differentially expressed (DE) genes. For DE analysis, counts were normalized using TMM method (built-in to edgeR). Any genes with less than in total 30 reads from all samples were dropped from further analysis. Differential expression (DE) analyses were conducted with the voom/limma analysis software packages by using total number of mapped reads as a scaling factor. Results were extracted from edgeR as tables and used for generating volcano or box plots in ggplot2 in RStudio.

To generate IGV genome browser tracks for ChIP- and RNA-seq data, all aligned bam files for each replicate of a given experiment were pooled and converted to BED format with bedtools bamtobed and filtered to be coverted into coverageBED format using the bedtools. Finally, bedGraphToBigWig (UCSC-tools) was used to generate the bigWIG files displayed on IGV browser tracks in the manuscript.

Gene ontology analysis was performed by using Statistical overrepresentation test and cellular component function PANTHER (http://pantherdb.org/). All genes being detected as expressed in RNA-sequencing data was used as reference. GO terms with at least 10 genes, at least 1.5-fold enrichment with less than 0.05 false discovery rate (FDR) were considered as significantly enriched. Significant GO terms were plotted in Prism 9.

### EEG recordings and behavioral monitoring

Implantations of Electroencephalograph (EEG) electrodes were performed in mice at age 12-16 weeks. EEG signals were recorded using two stainless steel screws inserted ipsilaterally into the skull. One was inserted 1.2 mm from the midline and 1.5 mm anterior to bregma, and the other was inserted 1.7 mm from midline and 2.25 mm posterior from to bregma. Seven days post-surgery, mice were transferred to individual behavior cages with a 12:12 h light/dark cycle and a constant temperature of about 23°C. Animals had access *ad libitum* to food and water and were allowed to recover from surgery for 7 days. Analysis was performed in individual cages equipped with overhead cameras (FLIR). Animals were connected to an amplifier (A-M Systems 1600) via a commutator. EEG signals were amplified and analog filtered (Gain 500, Low pass filter: 0.3 Hz, High pass filter: 100 Hz) and then digitized at 200Hz using Spike2 (CED Micro1401). Spontaneous sleep/wake behavior was monitored longitudinally via EEG recordings and video tracking. Epileptic episodes were scored manually by inspecting both the EEG signals and simultaneous video recordings.

### Quantification and statistical analysis

Analysis was conducted in R and with GraphPad Prism 9. Sample sizes were chosen based on previous experiments and literature surveys. No statistical methods were used to pre-determine sample sizes. Exclusion criteria used throughout this manuscript were pre-defined. See descriptions in the respective sections of the methods. Animals were randomly assigned to treatment groups. Appropriate statistical tests were chosen based on sample size and distribution of data points and are indicated in individual experiments.

