## Supplementary information for "Control of neuronal excitation-inhibition balance by BMP-SMAD1 signaling"

#### Extended Data Figures and Supplementary Information

Extended Data Fig. 1

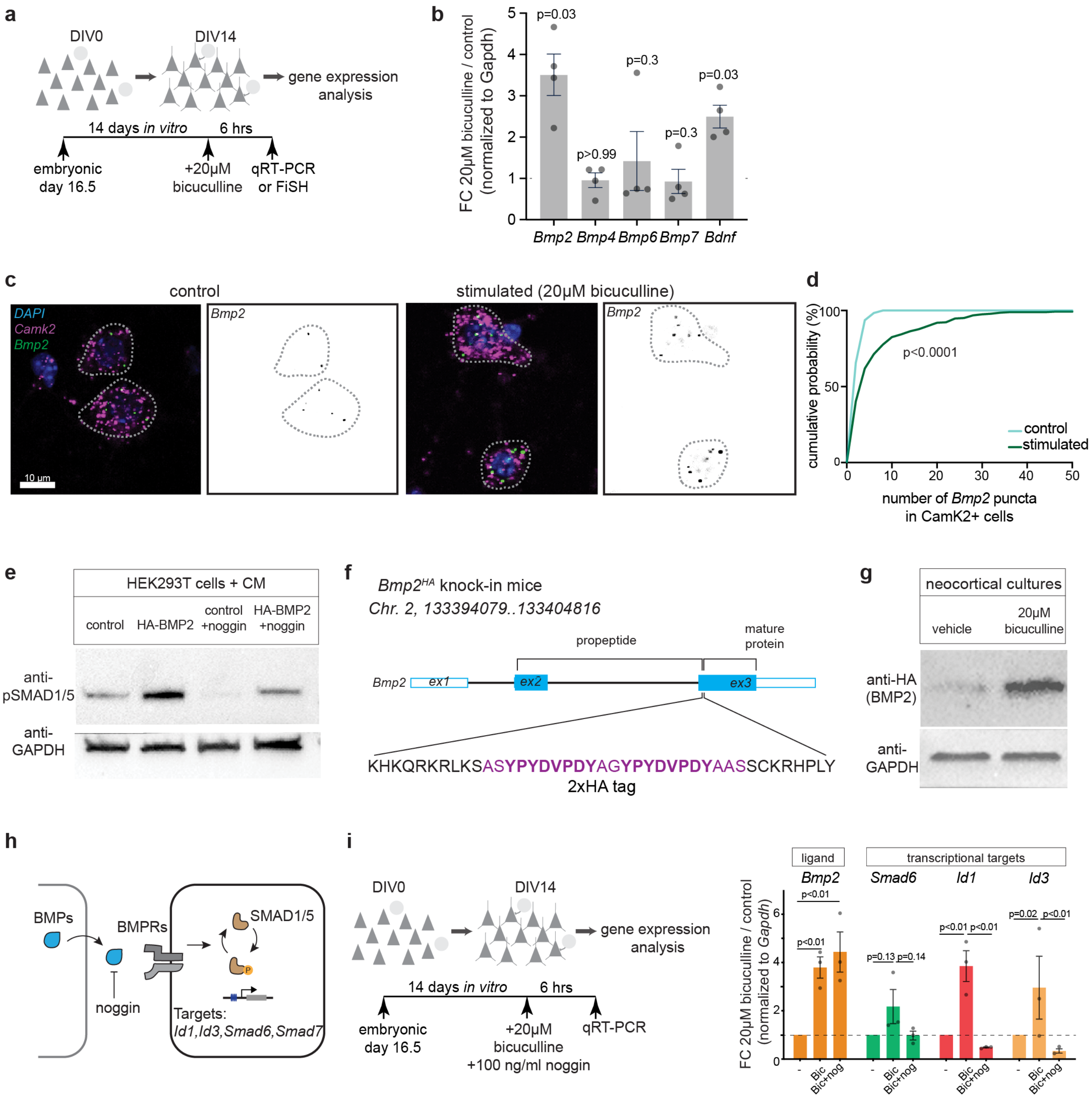


**Extended Data Fig. 1. Elevation of neuronal network activity triggers BMP2 upregulation in neocortical glutamatergic neurons *in vitro*. (a)** Schematic representation of cortical cultures and pharmacological activity manipulation. **(b)** qPCR assessment of *Bmp2*, *Bmp4*, *Bmp6*, Bmp7 and *Bdnf*

transcripts in DIV14 neocortical cultures treated with 20 μM bicuculline for 6 hours expressed as fold-change (FC) compared to untreated cultures. All expression values were normalized to *Gapdh* (N=4 independent cortical cultures, total of 3 technical replicates, Mann-Whittney test). **(c)** *Bmp2* transcript levels by fluorescence in situ hybridization (FiSH) in *Camk2*-positive glutamatergic neurons in naïve and stimulated neocortical cultures). **(d)** Cumulative distribution of *Bmp2* FiSH signal per cell in control and stimulated neurons (N=3 independent cortical cultures, Komolgorov-Smirnov test). **(e)** Confirming functional signaling for HA-epitope-tagged BMP2 in cultured cells. Western blot for phosphorylated SMAD1/5 (anti-pSMAD1/5) in cultured human embryonic kidney cells (HEK293T) treated with conditioned medium (CM) from control or HA-BMP2-expressing cells containing or lacking 100 ng/ml noggin. **(f)** Illustration of Crispr-based knock-in strategy for introduction of an epitope tag into the endogenous mouse *Bmp2* locus. A double HA tag sequence and flanking homology arms were encoded in a single stranded DNA oligo and were inserted in *Bmp2* exon 3 (ex3) at a Crispr/Cas9 cleavage site through homology-directed repair. The 2x HA tag is positioned at the N-terminus of the mature BMP2 protein. Resulting homozygous *Bmp2*$^{HA/HA}$ knock-in mice were viable and fertile. **(g)** Western blot with anti-HA antibodies of lysate from cultured neocortical neurons from *Bmp2*$^{HA/HA}$ knock-in mice (DIV14) either naïve or treated for 24 hours with 20μM bicuculline (N=3 independent cortical cultures). BMP2$^{HA}$ expression levels *in vivo* could not be reliably assessed, likely due to its low abundance in the complex tissue samples. **(h)** Illustration of inhibition of BMP-signaling by the extracellular antagonist noggin. **(i)** qPCR assessment of *Bmp2*, *Smad6*, *Id1* and *Id3* transcripts expressed as fold-change in bicuculline (bic, 20μM for 6 hours) and Bic+Nog (20μM bicuculline and 100 ng/ml noggin for 6 hours) compared to naïve cultures (N=3 independent cortical cultures, total of 3 technical replicates, two-way ANOVA).

Extended Data Fig. 2

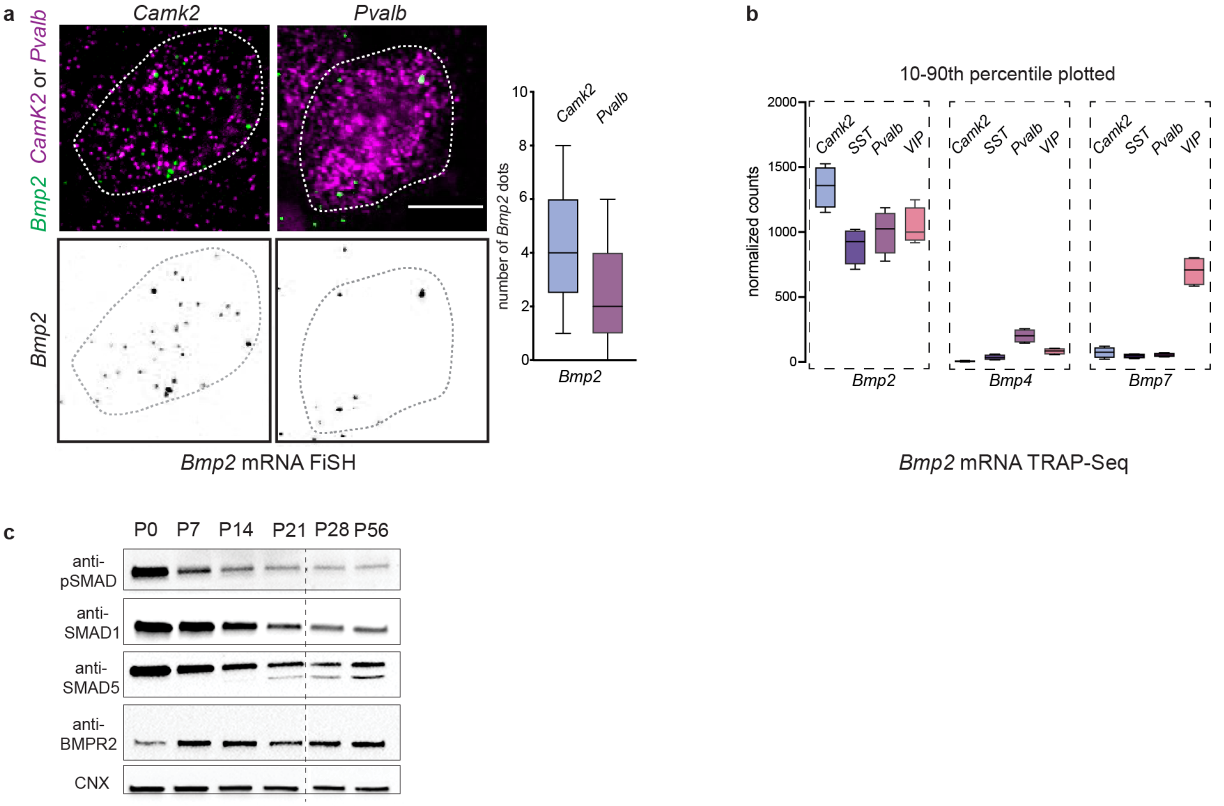


**Extended Data Fig. 2. Expression of BMP signaling components in the adult mouse neocortex.** **(a)** Quantification of *Bmp2* mRNA expression *Camk2*$^+$ and *Pvalb*$^+$ neurons in layer 2/3 of mouse barrel cortex (P25-30) assessed by FiSH (N=3 mice, n=57 cells/*Camk2*+ and n=45 cells/*Pvalb*+) (**b**) mRNA expression of *Bmp2*, *Bmp4*, *Bmp7* in P25 mouse neocortex in genetically-defined *Camk2*$^+$ principal neurons and somatostatin$^+$ (SST), PV, and Vasoactive intestinal peptide$^+$ (VIP) interneurons extracted from SPLICECODE database of TRAP-Seq analysis [72]. (**c**) Developmental expression levels assessed by Western blot of BMP receptor type 2 (BMPR2), transcriptional mediators (SMAD1 and SMAD5) and their active complex (pSMAD1/5/9) in the mouse neocortex (postnatal day 0 to postnatal day 56).

### Extended Data Fig. 3

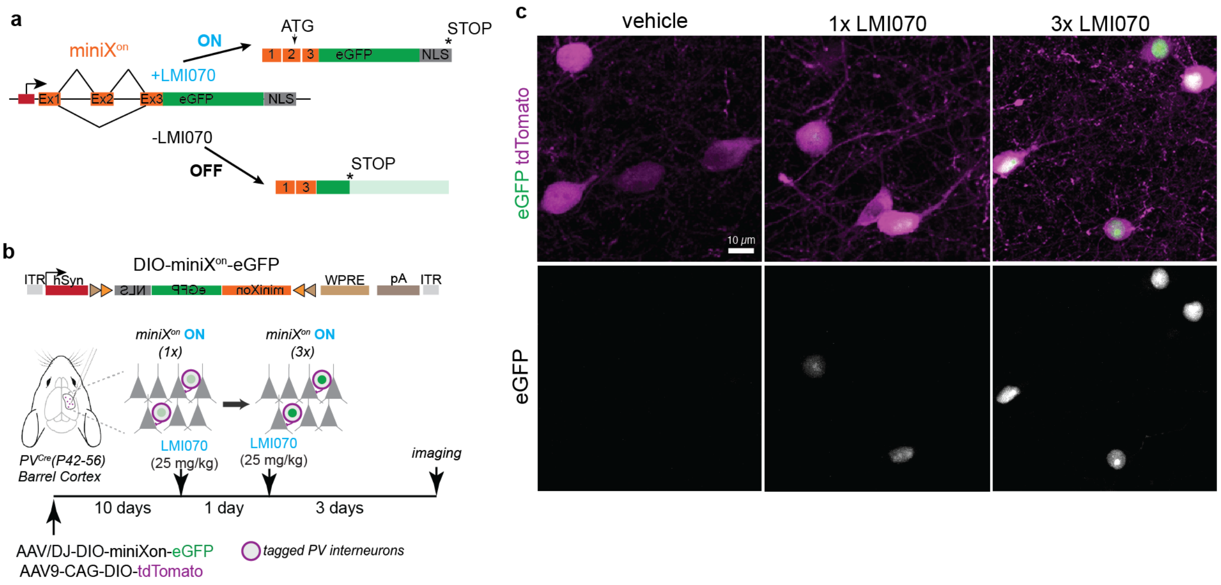


**Extended Data Fig. 3. Regulated expression from chemically-gated AAV X$^{on}$ in barrel cortex PV interneurons. (a)** Schematic illustration of miniX$^{on}$ regulation of protein expression[33]. In presence of the small molecule LMI070, alternative splicing of the cassette shifts to include a translational start codon in exon 2 (Ex2) and, thus, turns on expression of nuclear targeted eGFP reporter protein (NLS-eGFP). In the absence of LMI070, the AUG start codon-containing exon is skipped and translation does not occur in the correct reading frame. **(b)** Schematic diagram for cre-dependent expression of miniX$^{on}$ constructs in PV interneurons by AAV injection into the barrel cortex of adult *PV$^{cre}$* mice. **(c)** Representative images for nuclear NLS-eGFP expression in PV interneurons of mice treated by oral gavage with vehicle or 25 mg/kg LMI070 (1x or 3x in 24-hour intervals).

### Extended Data Fig. 4

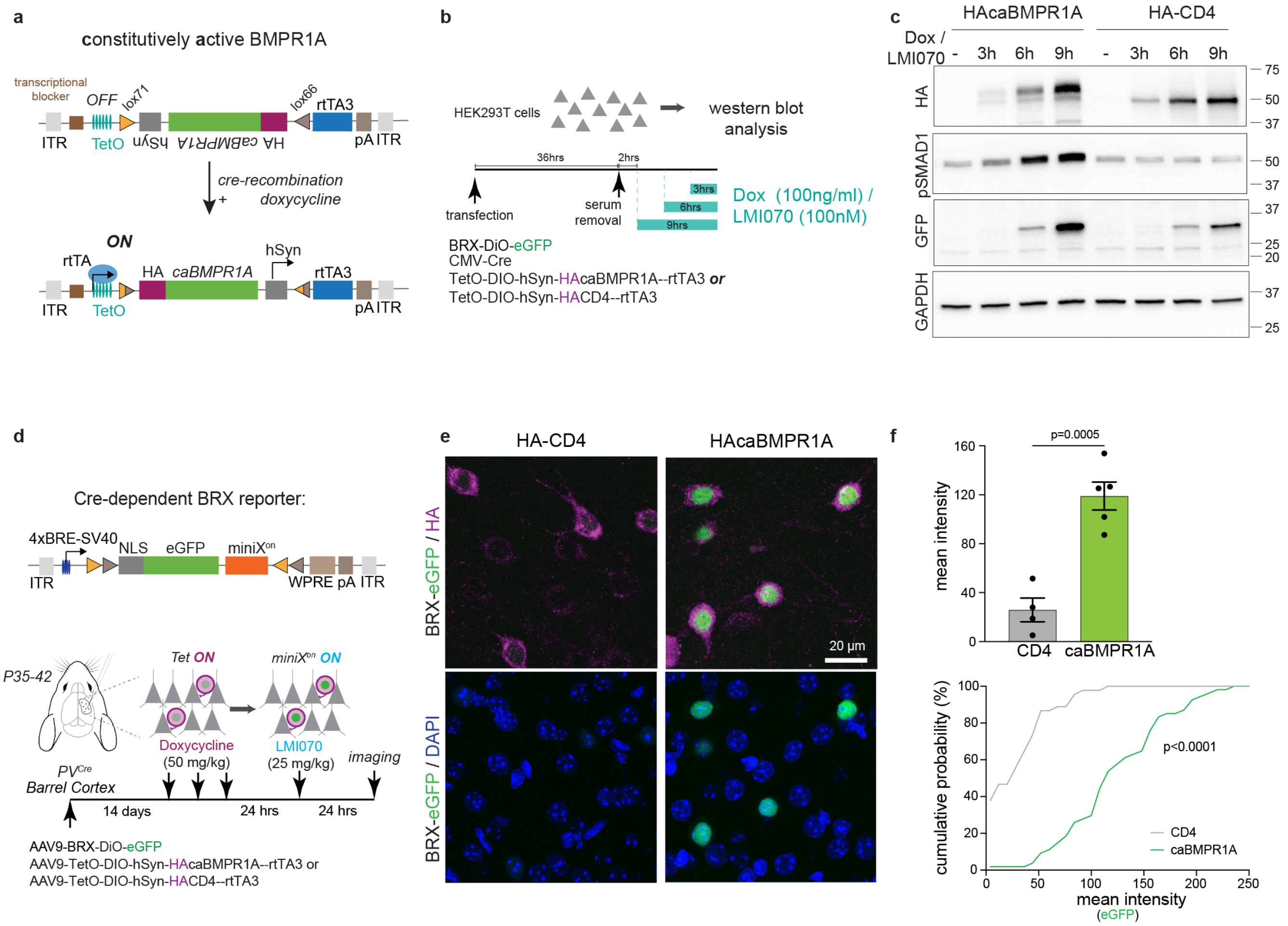


**Extended Data Fig. 4. Generation of a cre- and doxycycline-dependent expression vector for constitutively-active BMPR1A receptor. (a)** The vector contains a cre-dependent (lox71-lox66) inversion cassette encoding the open reading frame of the HA-epitope-tagged receptor (HA-caBMPR1A) and human synapsin promoter. Upon inversion, the reverse transcriptional activator rtTA3 is expressed from the synapsin promoter. In presence of doxycycline, rtTA3 drives transcription of HA-caBMPR1A from Tet operator sequences (TetO). (**b**) Experimental paradigm for testing regulated expression of HA-caBMPR1A and activation of the BRX-eGFP reporter in vitro. (**c**) Western blot analysis of cell lysates from transiently transfected HEK293 cells expressing BRX-DiO-eGFP, cre-recombinase under control of CMV promoter, and HA-caBMPR1A or a HA-CD4 control protein under control of the TetO elements. Upon chemical induction with doxycycline and LMI070, expression of HA-caBMPR1A results in an elevation of pSMAD1 signal and accumulation of eGFP from the BRX reporter. The low level of eGFP expression seen in the HA-CD4 control condition represents a low level of leakiness of expression from 4xBRE elements or activation due to endogenous BMP signaling events**. (d)** Vector for cre-dependent BRX-reporter where the BRX-NLS-eGFP cassette is inverted and flanked by loxP sites in a DiO configuration. Schematic diagram for cre-dependent co-expression of BRX-DiO-eGFP and HAcaBMPR1A constructs in PV interneurons by AAV injection into the barrel cortex of adult $PV^{cre}$ mice. **(e)** Representative images of BRX reporter signal in barrel cortex layer 2/3 of $PV^{Cre}$ mice. **(f)** Bar graph for mean ± SEM of nuclear eGFP intensity per mouse (N=4-5 mice/group, n=45-54 cells per condition, unpaired t-test) and cumulative distribution of eGFP reporter intensity per PV interneuron (Komolgorov-Smirnov test). Scale bar in e is 20 μm.

Extended Data Fig. 5

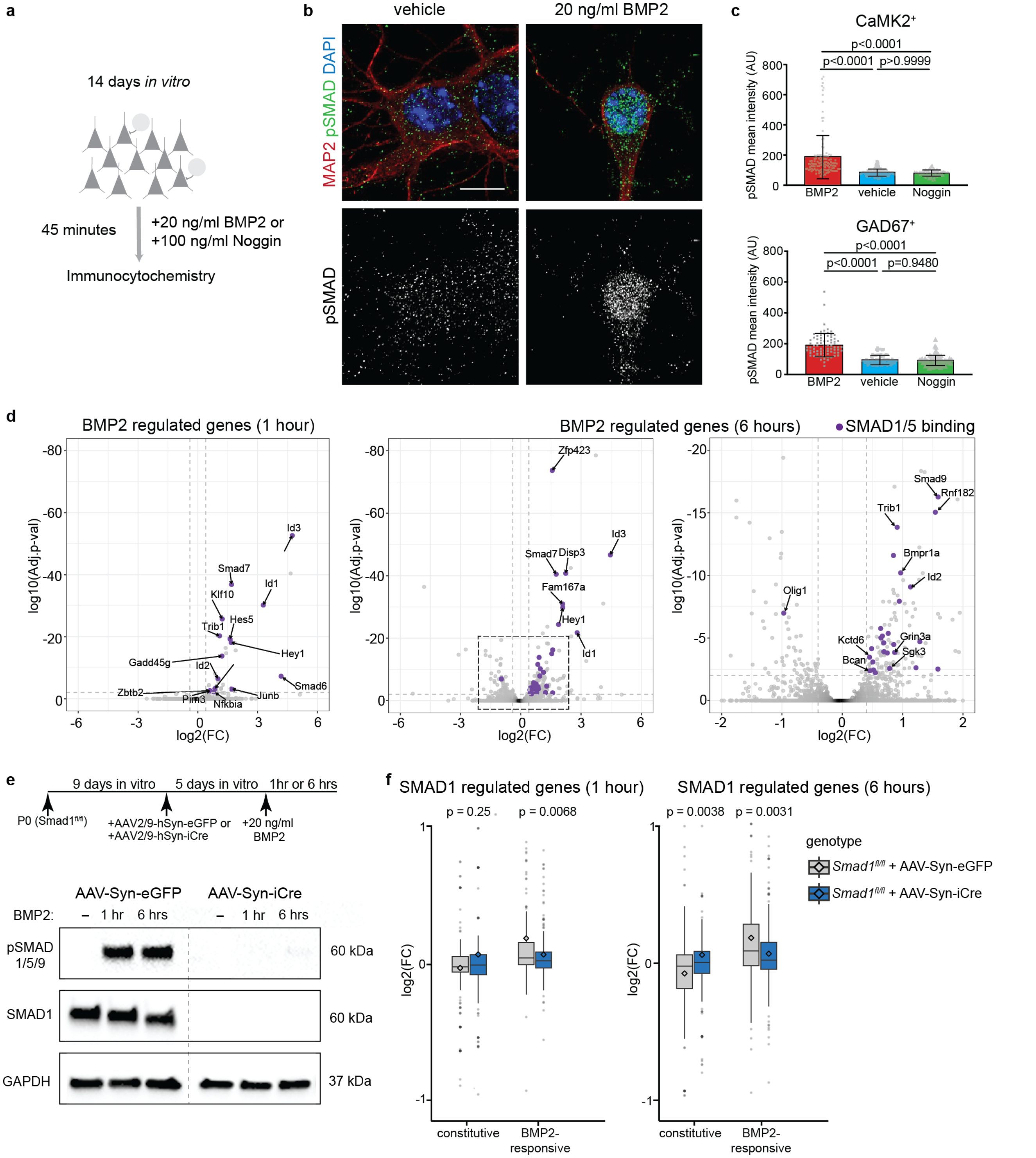


**Extended Data Fig. 5. BMP-SMAD1 signaling in neocortical neurons. (a)** Schematic representation of cortical cultures and BMP pathway manipulations. **(b)** Immunostaining of naïve or BMP2-stimulated (20ng/ml for 45 minutes) cultured neocortical neurons (DIV14) with antibodies to the neuronal marker microtubule associated protein-2 (MAP2) and pSMAD1/5/9 (activated SMAD). **(c)** Quantification of nuclear pSMAD intensity in cultured CaMK2$^{+}$ glutamatergic neurons and GAD67$^{+}$ GABAergic neurons from BMP2-treated (20ng/ml 45 minutes), vehicle-treated and noggin-treated (100 ng/ml, minutes) cortical cultures (N=3 independent cultures, one-way Anova followed by Tukey's multiple comparisons test, the bar graphs show then means ± SEM.) **(d)** Volcano plot of RNA-seq expression data from neocortical cultures (DIV14) stimulated with BMP2 (20ng/ml) for one hour (left) or 6 hours (middle and right). Log2 fold change (FC) of expression values for stimulated over non-stimulated cells and log10 adjusted p-values are displayed. Direct SMAD1/5 targets identified in ChIP-seq that are significantly regulated were marked in purple and are indicated by arrow. Gray dashed lines indicate 30% change and adj. p-value of 0.01 which were used as cut-offs to consider genes significantly regulated. Black dashed lines indicate the 2 fold change and log10 adj.p-values less than 20 which were used as cut-offs to highlight genes moderately but significantly changed (right). **(e)** Experimental design and Western blot for detection of SMAD1 and pSMAD1/5/9 protein levels in control (AAV-Syn-eGFP) and neuron-specific Smad1 conditional knock-out (AAV-Syn-iCre) cultured neocortical neurons (DIV14), either naïve (-) or treated with recombinant BMP2 (20 ng/ml) for 1 hr or 6 hrs (representative of N=3 independent cortical cultures). **(f)** Differential gene expression in neocortical cultures (DIV14) from *Smad1$^{fl/fl}$* mice infected with control protein (eGFP, displayed in gray) or cre recombinase (iCre, displayed in blue) expressing AAVs. The box plots show the log2 fold change in gene expression assessed by RNA-seq 1 hr (left) or 6 hrs (right) after BMP2 stimulation as compared to non-stimulated cultures. Genes with constitutive and BMP2-responsive SMAD1/5 binding events as identified by ChIP-seq are plotted separately. The statistically significant increase in expression of genes with constitutive SMAD1/5 binding events suggests that these genes are normally repressed by SMAD1. Horizontal black lines mark the median, whiskers indicate standard deviations and diamonds mark the mean of the fold changes (N=4 cultures/condition, p-values were obtained with Wilcoxon test).

#### Extended Data Fig. 6

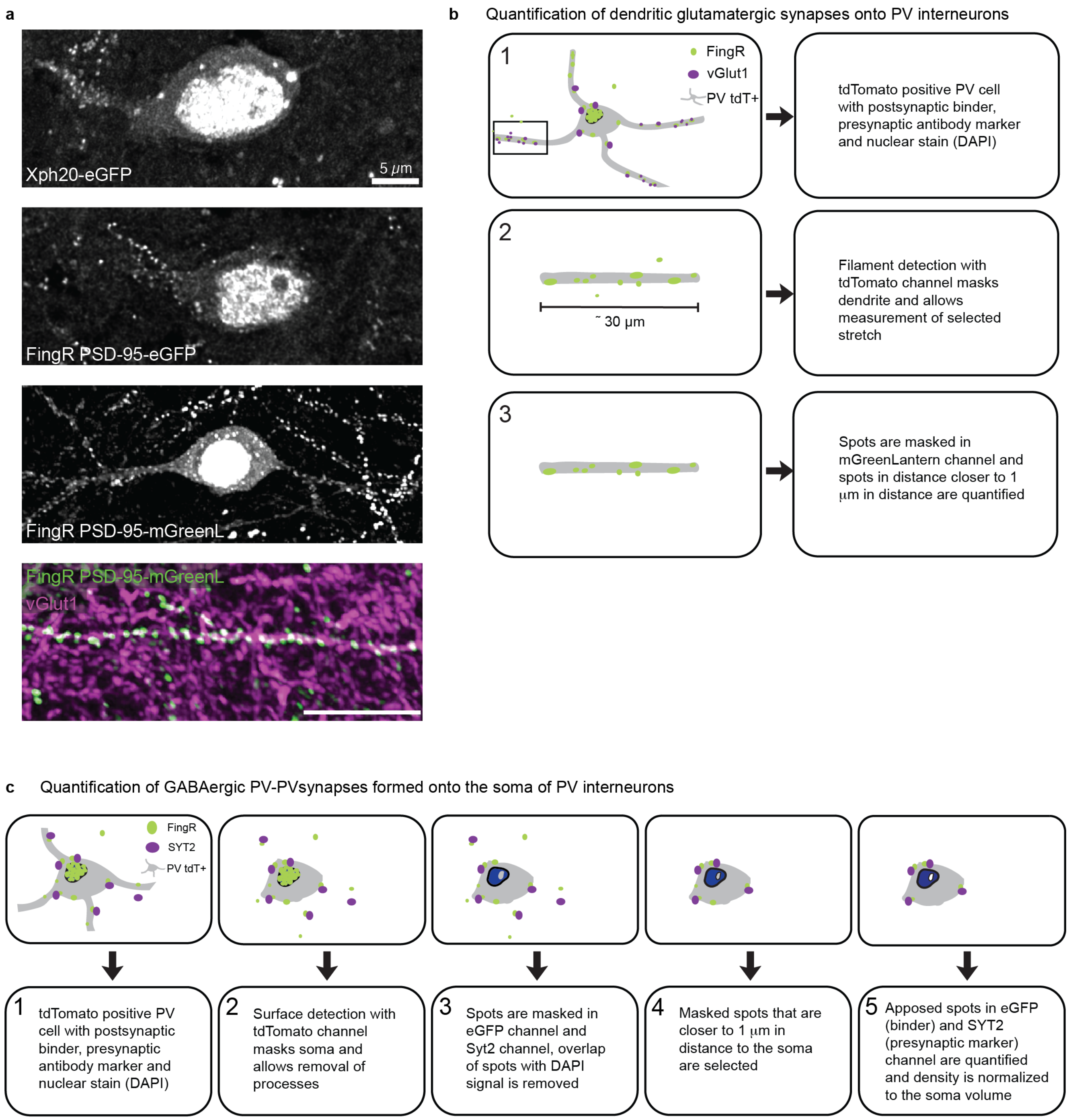


**Extended Data Fig. 6. Optimization of intrabody labelling and quantification of synaptic innervation of PV interneurons.** We optimized the original FingR-PSD-95-eGFP constructs [42] by introducing the neuron-optimized fluorophore mGreenLantern [73], and placing the cDNA under control of the neuron-specific synapsin promoter. We then compared FingR-PSD-95-mGreenLantern with FingR-PSD-95-eGFP and a PSD-95 paralog-specific Xph20-EGFP intrabody [74] by stereotaxic injection of cre-dependent AAVs into the barrel cortex of adult (P56-P72) *PV*$^{cre}$ mice. In these experiments, the FingR-PSD-95-mGreenLantern constructs yielded the most reproducible and discrete labeling of glutamatergic postsynaptic sites with little cytoplasmic or non-synaptic labeling. **(a)** Representative images of PV interneurons expressing Xph20-eGFP, PSD-95FingR-eGFP, PSD-95FingR-mGreenLantern. Scale bar in top panel is 5 μm. Co-immunostaining with the glutamatergic presynaptic marker VGlut1 (magenta) reveals extensive overlap with the postsynaptic FingR-PSD-95-mGreenLantern marker. **(b)** Illustration of 3D quantification protocol for glutamatergic synapses on PV interneuron dendrites with IMARIS software. **(c)** Quantification protocol developed in IMARIS to quantify peri-somatic GABAergic synapses labelled with FingRGPHN-eGFP intrabodies and co-stained with Syt2 on PV interneurons.

Extended Data Fig. 7

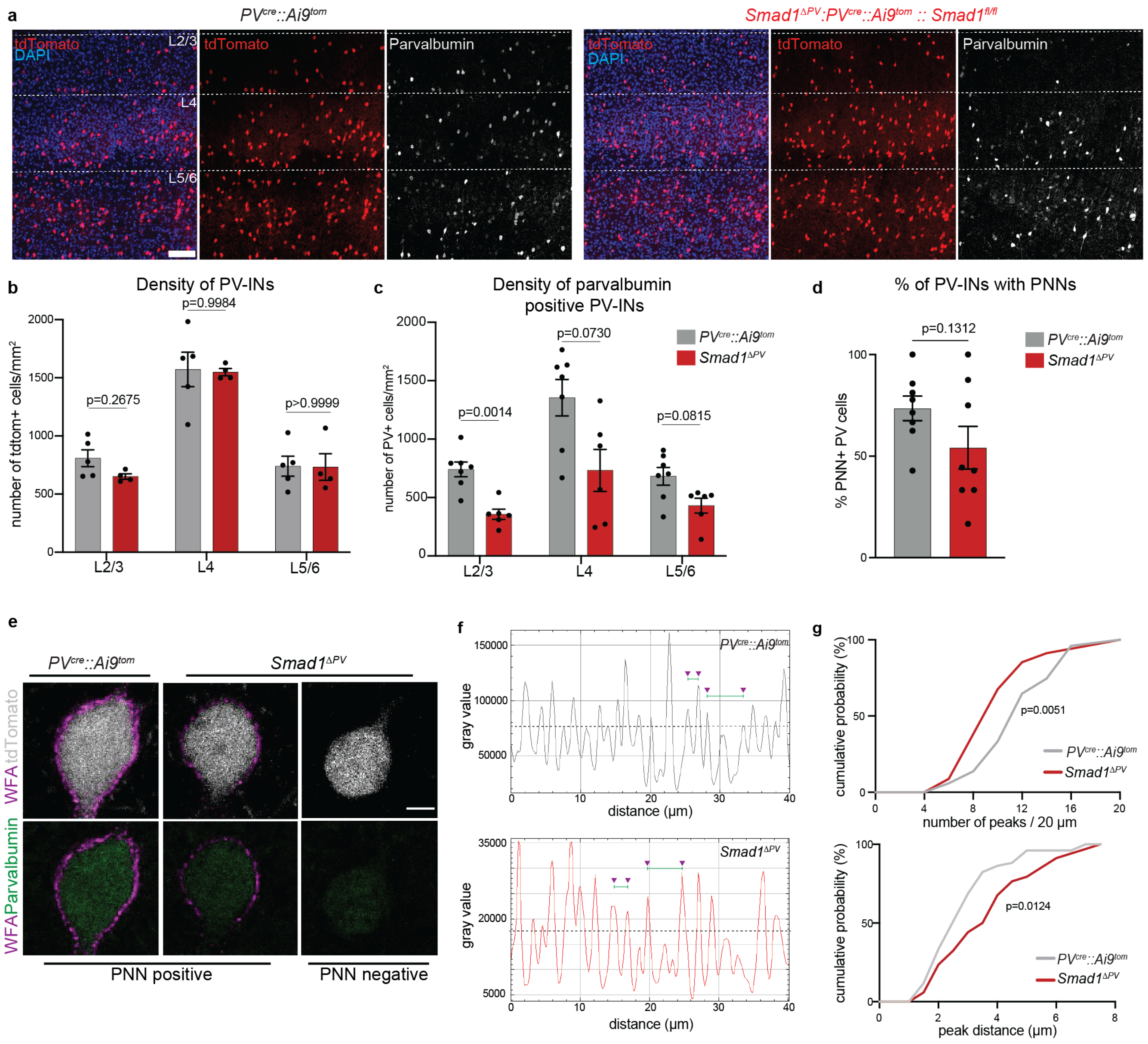


**Extended Data Fig. 7. Normal PV interneuron density but decreased parvalbumin protein immunoreactivity and reduced integrity of peri-neuronal nets in barrel cortex of *Smad1$^{\Delta PV}$* mice.** **(a)** Representative images of coronal sections of adult mouse (P56-P72) barrel cortex of *PV$^{cre}$::Ai9$^{tom}$* mice (left) and *Smad1$^{\Delta PV}$* mice (right) displaying nuclear DAPI, tdTomato, and anti-parvalbumin immunoreactivity. Scale bar is 100 μm **(b)** Quantification of density of tdTomato$^{+}$ PV interneurons across layers in barrel cortex of *PV$^{cre}$::Ai9$^{tom}$* mice (left) and *Smad1$^{\Delta PV}$* mice (N=4-5 mice/genotype, n=2 sections per genotype, mean cell density/mouse and SEM, two-way Anova followed with Sidak's multiple comparisons test). **(c)** Quantification of density of parvalbumin immunoreactive PV interneurons across layers in barrel cortex of *PV$^{cre}$::Ai9$^{tom}$* mice (left) and *Smad1$^{\Delta PV}$* mice (N=6-7 mice/genotype, n=2 sections per genotype, mean cell density/mouse and SEM, two-way ANOVA followed with Sidak's multiple comparisons test). (**d**) Quantification of PNN-positive and -negative cells (8 animals per genotype, mean and SEM, t-test. (**e**) High magnification views of individual PV interneurons in *PV*$^{cre}$*::Ai9*$^{tom}$ and *Smad1*$^{\Delta PV}$ mice, stained with WFA and anti-parvalbumin antibodies. For *Smad1*$^{\Delta PV}$ mice two examples are given – one PNN-positive and one PNN-negative cell. Scale bar is 5 μm. (**f**) Line graphs used for quantification of PNN integrity in PNN-positive cells from *PV*$^{cre}$*::Ai9*$^{tom}$ and *Smad1*$^{\Delta PV}$ mice. WFA staining intensity along a 20-40 μm line drawn at the center of the PNN structure is plotted. The number of peaks above a relative intensity threshold set at 50% of maximum peak height was quantified as well as inter-peak distances. For each plot 4 peaks and 2 examples of inter-peak intervals are marked. (g) Cumulative frequency plots for peak density and peak distance as described in f. 51- and 35-line graphs per genotype, derived from 8 animals per genotype. K.S. test.

#### Extended Data Fig. 8

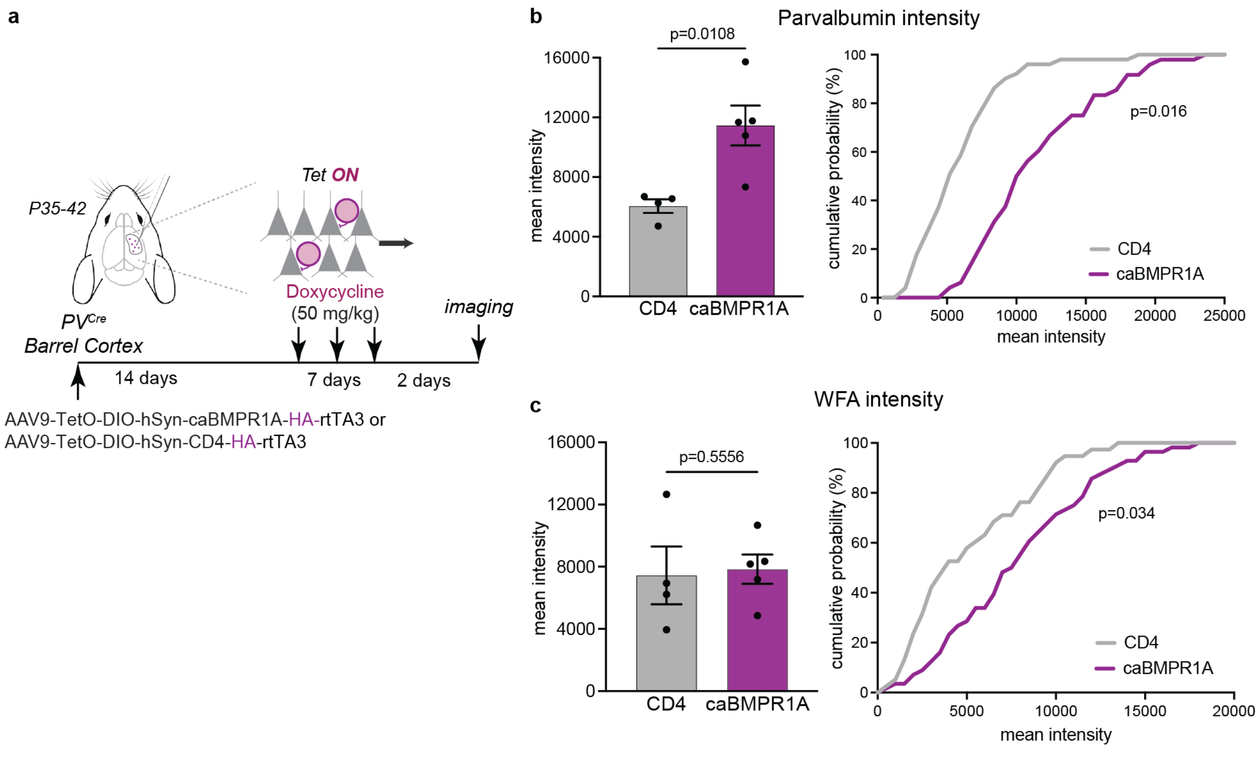


**Extended Data Fig. 8. Genetic activation of BMP-signaling in PV interneurons in adult somatosensory cortex results in an elevation of parvalbumin expression. (a)** Schematic diagram for expression of constitutively active BMPR1A (caBMPR1A) in PV interneurons. Recombinant AAVs encoding cre- and doxycycline-dependent expression cassettes (see Extended Data Figure 4 for details) were injected into the barrel cortex of adult *PV*$^{cre}$ mice. Expression of the caBMPR1A or the HA-CD4 control protein was induced by three intraperitoneal injections of doxycycline administered over one week and cells were analyzed 2 days after the final injection. **(b)** Mean intensity per animal and cumulative frequency of parvalbumin staining intensity in genetically-identified PV interneurons marked by the HA epitope on caBMPR1A or the CD4 control protein. **(c)** Mean intensity per animal and cumulative frequency of WFA staining intensity surrounding genetically-identified PV interneurons marked by the HA epitope on caBMPR1A or the CD4 control protein. N=4-5 animals, n= 48-51 cells.

#### Extended Data Fig. 9

**a**

| *PV interneurons (P56-72)* | *$PV^{cre}::Ai9^{tom}$* | *$Smad1^{\Delta PV}$* | *p-value* | *significance* |
|---|---|---|---|---|
| RMP (mV) | -75.25 ± 1.6 | -76.8 ± 1.7 | p=0.81 | ns |
| IR (MΩ) | 136.78 ± 7.1 | 113.0 ±8.4 | p=0.02 | * |
| Capacitance (pF) | 44.96 ± 3.7 | 48.61 ± 1.9 | p=0.19 | ns |
| AP-threshold (mV) | -40.1 ± 1.8 | -39.81 ± 1.0 | p=0.83 | ns |
| APmax (mV) | 45.15 ± 2.4 | 48.23 ± 2.0 | p=0.03 | * |
| AHP amplitude (mV) | 12.85 ± 0.5 | 16.74 ± 1.2 | p=0.02 | * |
| Half-width (ms) | 0.42 ± 0.03 | 0.47 ± 0.02 | p=0.49 | ns |
| MFF | 330.67 ± 22.4 | 256 ± 13.0 | p=0.04 | * |
| number of cells (n) | 12 | 14 | | |

**b**

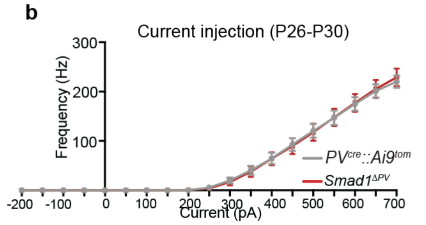


| *PV interneurons (P26-30)* | *$PV^{cre}::Ai9^{tom}$* | *$Smad1^{\Delta PV}$* | *p-value* | *significance* |
|---|---|---|---|---|
| RMP (mV) | -72.5 ± 1.0 | -71.0 ± 0.9 | p=0.16 | ns |
| IR (MΩ) | 99.2 ± 5.3 | 93.8 ± 5.1 | p=0.98 | ns |
| Capacitance (pF) | 37.1 ± 1.4 | 38.0 ± 1.8 | p=0.82 | ns |
| MFF | 288.3 ± 9.1 | 290 ± 11.7 | p=0.36 | ns |
| number of cells (n) | 23 | 21 | | |

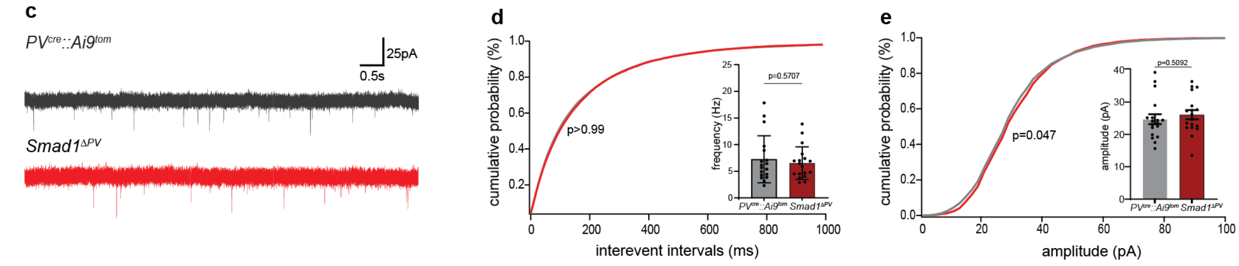


**Extended Data Fig. 9. (a)** Intrinsic and action potential properties of layer 2/3 PV interneurons from P56-72 *$PV^{cre}::Ai9^{tom}$* mice and *$Smad1^{\Delta PV}$* mice. RMP: resting membrane potential, IR: input resistance, MFF: maximum firing frequency, AP: action potential, AHP: afterhyperpolarization (N=4 mice, n=12 cells for *$PV^{cre}::Ai9^{tom}$* and N=4, n=14 cells for *$Smad1^{\Delta PV}$*, Kolmogorov-Smirnov test). **(b)** Comparison of firing frequencies of PV interneurons at given currents and their intrinsic properties from P26-30 *$PV^{cre}::Ai9^{tom}$* mice (gray) and *$Smad1^{\Delta PV}$* mice (red). RMP: resting membrane potential, IR: input resistance, MFF: maximum firing frequency (N=4 mice, n=23 cells for PVcre::Ai9tom and N=4 mice, n=21 cells for *$Smad1^{\Delta PV}$* mice, Kolmogorov-Smirnov test). (**c**) Representative traces of mEPSC recordings from control (gray) and *$Smad1^{\Delta PV}$* (red) PV interneurons in acute slice preparations from adolescent mice (P26-P30). **(d)** Frequency distribution of interevent intervals (Kolmogorov-Smirnov test) and mean mEPSC frequency (mean ± SEM for n=18 cells/genotype, from N=3 mice. Kolmogorov-Smirnov test).

**(e)** Frequency distribution of mEPSC amplitudes (Kolmogorov-Smirnov test) and mean mEPSC amplitude (mean ± SEM for n=18 cells/genotype, from N=3 mice. Kolmogorov-Smirnov test).

**Supplementary Movie 1. FingR-PSD-95 intrabody expression in *PV$^{Cre}$* mice.** Representative video generated from optical sections of FingR-PSD-95-mGreenLantern expressing layer2/3 PV interneuron showing the labelling of glutamatergic synapses forming on its dendrites and the soma. Images were taken from cleared 120-micron thick sections.

**Supplementary Movie 2. *Smad1$^{\Delta PV}$* mice have altered cortical network activity.** Representative EEG recording from *Smad1$^{\Delta PV}$* mice showing the brain activity of the mouse before, during and after a seizure event that occurred during a cage change.

**Supplementary Table 1.** Summary of ChIP-seq datasets collected from adult neocortex BMP2-stimulated primary cortical cultures.

**Supplementary Table 2.** Summary of RNA-seq datasets collected from BMP2-stimulated primary cortical cultures.

**Supplementary Table 3.** Summary of RNA-seq datasets collected from control and Smad1 mutant and BMP2 stimulated primary cortical cultures.

#### Additional Methods

**Recombinant DNA and viral vectors.** The following plasmids were obtained from Addgene: pAAV-hSyn-DIO-hM4D(Gi)-mCherry (RRID:Addgene_44362, gift from Bryan Roth), p4XBRE-SBE-SV40-GL3 (RRID:Addgene_67811, gift from Ron Prywes), pGL4.10/RSV_SF3B3-miniXon_Luciferase (RRID:Addgene_174660, gift from Beverly L. Davidson), FingR-PSD-95-eGFP-CCR5TC and FingR-Gephyrin-eGFP-CCR5TC (RRID:Addgene_46295 and RRID:Addgene_46296, gift from Donald Arnold), Xph20-eGFP-CCR5TC (RRID:Addgene_135530, gift from Daniel Choquet), pAAV-CAG-mGreenLantern (RRID:Addgene_164469, gift from Gregory Petsko), pAAV-S5E2-dTom-nlsdTom (RRID:Addgene_135630, gift from Jordane Dimidschstein).

pAAV-4xBRE-SV40-X$^{on}$-GFP and pAAV-4XBRE-SV40-DIO-X$^{on}$-GFP were generated by combining 4XBRE-SV40 promoter elements, the X$^{on}$ mini-cassette, and eGFP sequences, flanked by loxP and lox2272 sites in the case of double floxed inverted orientation (DIO). In pAAV-hSyn-DIO-X$^{on}$-GFP the 4XBRE-SV40 sequence was replaced by the human Synapsin (hSyn) promoter. pAAV-Tet-DIO-hSyn-HAcaBMPR1A-rtTA3 and pAAV-Tet-DIO-hSyn-HAcaCD4-rtTA3 were generated by combining human BMPR1A sequence with Q233D mutation or human CD4 sequence, hSyn promoter, Tet-operon casette, doxycycline- inducible rtTA3 transcriptional activator, HA tag sequence, and flanked by lox66 and lox71 sites for double floxed inverted orientation (DIO).

For conditional viral expression of intrabodies in PV interneurons, FingR or Xph20 coding sequences were introduced into pAAV vectors in double-floxed inverted orientation under control of the hSyn promoter combined with the CCR5 zinc finger interaction site. For expression of FingR-PSD-95 binder selectively but in a cre-independent manner in PV interneurons, S5E2 enhancer sequence was introduced into pAAV vectors upstream of FingR coding sequence combined with CCR5 zinc finger interaction site. For both enhanced conditional and S5E2-PSD-95 binders, eGFP was replaced by mGreenLantern coding sequence.

Viral supernatants were produced by co-transfection of HEK293T cells grown on 15cm dishes using calcium phosphate transfection of 70µg of AAV helper plasmid (Rep/Cap, Serotype 9), 200µg of AAV pHGTI-adeno1 (Plasmid factory) and 70µg of AAV vector plasmid carrying the cDNAs to be expressed. 45-60h after transfection, medium containing viral particles was harvested and purified using the Iodixanol purification method. Viral preparations were concentrated in Millipore Amicon 100K columns at 4°C. Virus samples were suspended in PBS, frozen in aliquots and stored at -80°C. Viral titers were determined by qPCR and were >$10^{13}$ particles/mL.

**Primary neuron culture.** Cortical cultures were prepared from E16.5 mouse embryos or newborn (P0) mice. Neocortices were digested by addition of papain (145 units in 7 ml, Worthington Biochemical #LK003176) for 30 min at 37°C and then mechanically dissociated. Cells were maintained in neurobasal medium (Gibco #21103) containing 2% B27 supplement (Gibco #17504-044), 2mM Glutamax (Gibco #35050-038), and 1% penicillin/streptomycin (Gibco, #15140122) at 37°C / 5% $CO_2$. For neuronal network activity stimulation experiments, cortical cultures were treated for 6 hours at day *in vitro* 14 (DIV14) with 25 mM KCl or with 20 µM bicuculline (Tocris #0130). For immunohistochemistry, ChiP-seq or RNA-seq experiments, DIV14 cultures were stimulated with 20 ng/ml human recombinant BMP2 (R&D systems, #355-BM-050), recombinant mouse Noggin (R&D Systems, 1967-NG-025), or vehicle. For neuron-specific *Smad1* loss of function experiments, cortical cultures from P0 *Smad1*$^{fl/fl}$ mice were infected at DIV9 with AAV9-hSyn-iCre virus at a multiplicity of infection of 20,000 or with a AAV9-hSyn-eGFP virus as negative control.

**RNA isolation and reverse transcription.** Mice were anesthetized with isoflurane (Baxter AG, Vienna, Austria) and brain was taken out into ice-cold PBS solution. Injected area was then dissected under Binocular Stereo Microscope (Olympus #MVX10) by using red fluorescence signal from mCherry expression. Dissected tissue was harvested in Trizol reagent (Sigma, T9424). Total RNAs were isolated and DNase treated on columns (RNeasy Micro kit, Qiagen, 74004) following the manufacturer's instructions. The cDNA libraries were built using between 100 and 200ng RNA reverse transcribed with ImPromII Reverse Transcriptase (Promega, #M314A), RNasin™ Plus RNase Inhibitor (Promega, #N261B), ImPromII 5X Reaction Buffer (Promega, #M289A), dNTPs (Sigma, D7295) and oligo$(dT)_{15}$ primer (Promega, C1101). Primary cortical cultures were washed 1x with PBS and lysed using Trizol reagent (Sigma, T9424) and followed by total RNA isolation and cDNA library preparation as described above.

Real-time quantitative PCRs were performed either with FastStart Universal SYBR GreenMaster (Roche, 04-913-850-001) or FastStart Universal Probe Master (Roche, 04-913-195-7001). PCRs were carried out in a StepOnePlus qPCR system (Applied Biosystems) and were analyzed with the StepOne software. Gene expression assays were used either with FastStart Universal SYBR GreenMaster (Roche, 04-913-850-001) or TaqMan Master Mix (Applied Biosystems) and comparative $C_T$ method. The mRNA levels were normalized to housekeeping β-actin mRNA or to Gapdh mRNA. For each assay, two to three technical replicates were performed and the mean was calculated.

Commercially available gene expression assays for Fos (Mm00487425_m1), Bdnf (Mm04230607_s1), *Id1 (Mm00775963_g1), Id3* (Mm01188138_g1)*, Smad6* (Mm00484738_m1)*, Smad7* (Mm00484742_m1)*, ActB (Mm00607939_s1)* (Mm.PT.58.13518911), *Bmp2*

(Mm01340178_m1), *Bmp4* (Mm00432087_m1), *Bmp6* (Mm01332882_m1), *Bmp7* (Mm00432102_m1), *Gapdh* (Mm99999915_g1) were from ThermoFisher.

Custom primer sequences were as follows:

| Primer Name | Sequence |
|---|---|
| Brevican-Exon11-Fwd | 5'-CAT CGA GGG TGA CTT CTT GT-3' |
| Brevican-Exon12-Rev | 5'-ACC ATG ACC ACA CAG TTC TC-3' |
| Id3-Exon1-Fwd | 5'-GCA GCG TGT CAT AGA CTA CAT C-3' |
| Id3-Exon2-Rev | 5'-GTC CTT GGA GAT CAC AAG TTC C-3' |
| Grin3a-Exon7-Fwd | 5'-CTG CTG CTA CCA CGA ATC AA-3' |
| Grin3a-Exon8-Rev | 5'-TCT TGG AAC ATG GCT GCT T-3' |
| ActB-Exon5-Fwd | 5'-AGA TTA CTG CTC TGG CTC CTA-3' |
| ActB-Exon6-Rev | 5'-CTG CTT GCT GAT CCA CAT CT-3' |

**Western blotting.** Primary cortical cells were lysed in 50 mM Tris HCl pH 7.5, 150 mM NaCl, 10% Glycerol, protease inhibitor Roche Complete™ mini, 1% Triton X-100). Transfected HEK293T cells were lysed in 50 mM Tris HCl, 150 mM NaCl, 1.0% (v/v) NP-40, 0.5% (w/v) Sodium Deoxycholate, 1.0 mM EDTA, 0.1% (w/v) SDS). Lysates were centrifuged for 10 minutes at 16'000 g at 4°C and solubilized proteins were analyzed by polyacrylamide gel electrophoresis on 4%-20% gradient gels (BioRad, 4561093) followed by transfer onto nitrocellulose membrane. For enhanced chemiluminescence detection, WesternBright ECL kit (Advansta #K 12045-D20) and WesternBright Quantum (Advansta #K-12042-D20) were used. Signals were acquired using an image analyzer (Bio-Rad, ChemiDoc MP Imaging System and Li-Cor, Odyssey).

**Fluorescent *in situ* hybridization.** Multiplex fluorescent *in situ* hybridization (FiSH) was performed using the RNAScope Fluorescent Multiplex Kit (Advanced Cell Diagnostics, ACD). Mouse brains were snap frozen in liquid nitrogen and 18 µm coronal sections were cut between Bregma -1.43 and -2.15 (including barrel cortex and dorsal hippocampus) on a cryostat. Sections were fixed at 4°C overnight with 4% paraformaldehyde in 100mM phosphate buffered saline, pH 7.4. The procedure followed the manufacturers' instructions. For *in vitro* FiSH experiments, cortical cultures were plated onto glass coverslips. At DIV12, cells were fixed with 4% PFA for 15 minutes. The following probes were used: *Bmp2* (ACD #406661), *Camk2* (ACD #411851) and *Pvalb* (ACD #421931). Images were acquired with an upright LSM700 confocal microscope (Zeiss) using 40x/1.3 or 63x/1.4 Apochromat objectives. Cell types were identified based on the presence of the corresponding marker transcript. A region of interest (ROI) was drawn to define the area of the cell and dots in the ROI were manually counted. The number of dots in the ROI were then normalized to the cell area. Image acquisition and counting was done blinded to the experimental condition.

**Electrophysiology.** Cortical slice preparation from adolescent (P26-P28) or adult mice (P56-72) was adapted from previously described protocols [14]. Briefly, animals were anesthetized with isoflurane (Baxter AG, Vienna, Austria). Parasaggital slices of 300 µm were cut with a vibratome (VT1200S, Leica) in ice-cold oxygenated (95% $O_2$/5% $CO_2$) NMDG solution (93 mM NMDG, 93 mM HCl, 2.5 mM KCl, 1.2 mM $NaH_2PO_4$, 30 mM $NaHCO_3$, 20 mM HEPES, 25 mM glucose, 5 mM sodium ascorbate, 2 mM

Thiourea, 3 mM sodium pyruvate, 12 mM N-acetyl L-cysteine, 10mM $MgSO_4$ and 0.5 mM $CaCl_2$, pH 7.35). Slices were kept at 33.0 ± 1 °C in oxygenated NMDG solution for 12 minutes and then transferred to artificial cerebrospinal fluid (aCSF; 125 mM NaCl, 2.5 mM KCl, 1.25 mM $NaH_2PO_4$, 24 mM $NaHCO_3$, Na-Ascorbate (5 mM), 12.5 mM glucose, 1 mM $MgCl_2$ and 2 mM $CaCl_2$, pH 7.4) and kept at room temperature for at least 1 h before starting the recordings. During the recording sessions the slices were held in a custom chamber heated to 33.0 ± 1 °C with oxygenated aCSF perfusion.
Whole-cell patch-clamp recordings from layer2/3 PV interneurons of the barrel cortex were performed in voltage or current clamp mode using a Multiclamp 700B amplifier (Molecular Devices, Sunnyvale, CA) and under visualization in an upright microscope (Olympus) equipped with gradient contrast infrared visualization (Luigs and Neumann) using a 60× objective. For all experiments, data was digitized by Digidata 1440a (Molecular Devices) at 10 kHz and filtered at 1 kHz. For mEPSC and mIPSC recordings, patch pipettes (3–6 MΩ) were pulled with Sutter P-1000 micropipette puller (Sutter Instruments) and were filled with the following intracellular solution: 130 mM $CsMeSO_3$, 8 mM NaCl, 4 mM Mg-ATP, 0.3 mM Na-GTP, 0.5 mM EGTA, 10 mM HEPES, 5 mM QX314. For excitability measurements, intracellular solution composition is 142 mM K-gluconate, 10 mM HEPES, 1 mM EGTA, 2.5 mM $MgCl_2$, 4 mM Mg-ATP, 0.3 mM Na-GTP, 10 mM Na-phosphocreatine. mEPSCs and mIPSCs were recorded in the presence of 1 μM tetrodotoxin (TTX). To isolate both mEPSCs and mIPSCs from the same cell, mEPSCs were recorded at -70 mV holding potential and mIPSCs at 0 mV holding potential. Cells with >20% change in the series resistance were excluded from the analysis. mEPSCs and mIPSCs were analyzed using a template-matching algorithm implemented in Clampfit 10 (Molecular Devices). Automatically detected events were visually controlled and false positive events were deleted. Amplitude and frequencies were then manually analyzed. Input resistance, membrane time constant and capacitance were also calculated in Clampfit 10. Action potentials of PV interneurons were automatically detected with Neuromatic [15] and analyzed using a custom-made script written for this project in Igor Pro 8 software (WaveMetrics).

Cortical slice preparations and excitability measurements from layer 2/3 PV interneurons of P26-P30 mice were performed as described above with the following modifications: Slices were cut in sucrose-based solution: 75 mM sucrose, 87 mM NaCl, 25 mM $NaHCO_3$, 2.5 mM KCl, 1.25 mM $NaH_2PO_4$, 0.5 mM $CaCl_2$, 7 mM $MgCl_2$ and 10 mM glucose. Slices were immediately transferred to a storage chamber containing artificial cerebral spinal fluid (aCSF) containing: 125 mM NaCl, 25 mM $NaHCO_3$, 2.5 mM KCl, 1.25 mM $NaH_2PO_4$, 2 mM $MgCl_2$, 2.5 mM $CaCl_2$ and 11mM glucose, pH 7.4, constantly bubbled with 95% O2 and 5% CO2; 315-320 mOsm. Slices were maintained at 35°C in aCSF for 60 min and then kept at room temperature before their transfer to the recording chamber. During the recordings, the slices were continuously perfused with aCSF at 35.0 ± 2.0°C throughout the experiments. Neuronal activity of layer 2/3 PV interneurons was recorded with borosilicate glass pipettes (4-6 MΩ) filled with an intracellular solution containing: 125 mM K-gluconate, 20 mM KCl, 10 mM HEPES, 10 mM EGTA, 2 mM $MgCl_2$, 2 mM $Na_2ATP$, 1 mM $Na_2$-phosphocreatine, 0.3 mM $Na_3GTP$ and 0.2% biocytin. Passive membrane properties and detection of action potentials were measured by using a custom-made script in Igor Pro 8 software (WaveMetrics).

**Open field test.** Behavioral testing was done with males and females which were aged between 10 and 16 weeks. Mice were handled for at least 3 days before the test and acclimatized to the testing room for at least an hour before starting the experimentation. Mice were placed in the center of a rectangular OFT box (30 cm in width and 45 cm in length, with 30-cm-tall walls) for 15 minutes. Videos were recorded with a downward-facing camera from above with ANY-maze at a rate of up to 30 Hz. Distance (as cm) was extracted from ANY-MAZE and velocity (cm/min) was calculated.

#### Methods References


1 Furlanis, E., Traunmuller, L., Fucile, G. & Scheiffele, P. Landscape of ribosome-engaged transcript isoforms reveals extensive neuronal-cell-class-specific alternative splicing programs. *Nature neuroscience* **22**, 1709-1717 (2019). https://doi.org:10.1038/s41593-019-0465-5

2 Monteys, A. M. *et al.* Regulated control of gene therapies by drug-induced splicing. *Nature* **596**, 291-295 (2021). https://doi.org:10.1038/s41586-021-03770-2

3 Gross, G. G. *et al.* Recombinant probes for visualizing endogenous synaptic proteins in living neurons. *Neuron* **78**, 971-985 (2013). https://doi.org:10.1016/j.neuron.2013.04.017

4 Campbell, B. C. *et al.* mGreenLantern: a bright monomeric fluorescent protein with rapid expression and cell filling properties for neuronal imaging. *Proc Natl Acad Sci U S A* **117**, 30710-30721 (2020). https://doi.org:10.1073/pnas.2000942117

5 Rimbault, C. *et al.* Engineering paralog-specific PSD-95 synthetic binders as potent and minimally invasive imaging probes. *bioRxiv*, 2021.2004.2007.438431 (2021). https://doi.org:10.1101/2021.04.07.438431

6 Huang, S. *et al.* Conditional knockout of the Smad1 gene. *Genesis* **32**, 76-79 (2002). https://doi.org:10.1002/gene.10059

7 Hippenmeyer, S. *et al.* A developmental switch in the response of DRG neurons to ETS transcription factor signaling. *PLoS Biol* **3**, e159 (2005). https://doi.org:10.1371/journal.pbio.0030159

8 Madisen, L. *et al.* A robust and high-throughput Cre reporting and characterization system for the whole mouse brain. *Nature neuroscience* **13**, 133-140 (2010). https://doi.org:10.1038/nn.2467

9 Richardson, C. D., Ray, G. J., DeWitt, M. A., Curie, G. L. & Corn, J. E. Enhancing homology-directed genome editing by catalytically active and inactive CRISPR-Cas9 using asymmetric donor DNA. *Nat Biotechnol* **34**, 339-344 (2016). https://doi.org:10.1038/nbt.3481

10 Rosenbloom, K. R. *et al.* The UCSC Genome Browser database: 2015 update. *Nucleic acids research* **43**, D670-681 (2015). https://doi.org:10.1093/nar/gku1177

11 Gaidatzis, D., Lerch, A., Hahne, F. & Stadler, M. B. QuasR: quantification and annotation of short reads in R. *Bioinformatics* **31**, 1130-1132 (2015). https://doi.org:10.1093/bioinformatics/btu781

12 Heinz, S. *et al.* Simple combinations of lineage-determining transcription factors prime cis-regulatory elements required for macrophage and B cell identities. *Mol Cell* **38**, 576-589 (2010). https://doi.org:10.1016/j.molcel.2010.05.004

13 Schindelin, J. *et al.* Fiji: an open-source platform for biological-image analysis. *Nat Methods* **9**, 676-682 (2012). https://doi.org:10.1038/nmeth.2019

14 Jiang, X. *et al.* Principles of connectivity among morphologically defined cell types in adult neocortex. *Science (New York, N.Y* **350**, aac9462 (2015). https://doi.org:10.1126/science.aac9462

15 Rothman, J. S. & Silver, R. A. NeuroMatic: An Integrated Open-Source Software Toolkit for Acquisition, Analysis and Simulation of Electrophysiological Data. *Front Neuroinform* **12**, 14 (2018). https://doi.org:10.3389/fninf.2018.00014
